# Neurons have an inherent capability to learn order relations: A theoretical foundation that explains numerous experimental data

**DOI:** 10.1101/2025.03.17.642834

**Authors:** Yukun Yang, Wolfgang Maass

**Author notes:** Contributing authors.

## Abstract

Brains are able to extract diverse relations between objects or concepts, and to integrate relational information into cognitive maps that form a basis for decision making. But is has remained a mystery how relational information is learnt and represented by neural networks of the brain. We show that a simple synaptic plasticity rule enables neurons to absorb and represent relational information, and to use it for abstract inference. This inherent capability of neurons and synaptic plasticity is supported by a rigorous mathematical theory. It explains experimental data from the human brain on the emergence of cognitive maps after learning several linear orders, it explains the terminal item effect that enhances transitive inference if a terminal item of an order is involved, and it provides a simple model for fast configuration of internal order representations in the face of new evidence. We also present a rigorous theoretical explanation for the surprising fact that 2D projections of neural representations of linear orders are curved, rather than linear. Since our model does not require stochastic gradient descent in deep neural networks for learning order relations, it is suited for porting the capability to learn multiple relations and using them for fast inference into edge devices with a low energy budget.

## 1 Introduction

A hallmark of higher cognitive function is the capability to extract and represent information about relations between objects in a visual scenes, or between abstract concept, to integrate information from several learnt relationships, and to use it for fast inference. In particular, the brain is able to extract information about order relations between items. Furthermore, it encodes this order information in a way that supports transitive inference, a key paradigm for abstract reasoning. Essential for transitive inference is the extraction and neural representation of a latent variable from items in an order relation that can not be directly observed: their rank. Experimental data show that brains are able to extract this latent variable from dispersed information about order relations between pairs of items, and to encode it in the parietal cortex. In fact, experimental show that rank information is encoded by the brain in two different ways: By a summation code where many neurons fire with a rate that is indicative of the rank of an item, and by neurons that only fire for a single rank and a few adjacent ranks (heterogeneous code), see the review in (Summerfield et al., 2020). Experimental evidence for the existence of a summation code was provided by (Ganguli et al., 2008; Fitzgerald et al., 2013; Rutishauser et al., 2018; Chafee, 2013), see section 6 of the review (Summerfield et al., 2020) and Box 3 in (Bottini and Doeller, 2020) for an overview. The summation code was explained in (Ganguli et al., 2008) from a more general perspective: The surprising dominance of a single dimension in the network dynamics of area LIP (lateral intraparietal cortex). This effect is consistent with a large body of experimental data on the representation of numbers and other types of magnitudes in the brain (Nieder and Dehaene, 2009; Roitman et al., 2012; Nieder, 2016; Lyons et al., 2016; Lorenzi et al., 2021; Luyckx et al., 2019; Summerfield et al., 2020; Bottini and Doeller, 2020; Viganó et al., 2024; Xu et al., 2024). Informally one refers to these common types of 1-D neural representations as mental number line and as a 1-D cognitive map (Bottini and Doeller, 2020). It is closely related to the theoretically postulated geometric mental line of (Di Antonio et al., 2024). One functional advantage of such a scalar neural network code is that downstream network can use the same decoder for many tasks (Fitzgerald et al., 2013).

The presence of a summation code for ranks of items in an order makes transitive inference easy: Downstream networks just have to compare the firing activity of these neurons for two different items in an order. In fact, this computational strategy automatically gives rise to the symbolic distance effect that is seen in human behavior, see the review in (Mannella and Pezzulo, 2024): Human subjects have lower reaction time and make fewer errors if the required transitive inference involves pairs of items whose ranks are substantially different. But a summation code does not explain the terminal item effect, also referred to as serial position effect (Merritt and Terrace, 2011; Brunamonti et al., 2016; Mannella and Pezzulo, 2024), whereby transitive inference that involves a terminal item in an order exhibits fewer errors and is carried out faster than for other pairs of items with the same rank difference.

It remains unknown by which learning process the underlying latent variable of an order, the rank of an item, is extracted from dispersed pieces of information about order relations between pairs of items and mapped to the largely 1-dimensional dynamics of area LIP. Previous models for learning order relations in the brain involved multi-layer perceptrons (Nelli et al., 2023) or recurrent neural networks (Lippl et al., 2024) that were trained through stochastic gradient descent, or involved a special-purpose neuromodulatory system that had been optimized for this task (Miconi and Kay, 2025). An exception is the approach of (Di Antonio et al., 2024), where also a shallow linear network is used for learning orders. They considered the case where all order relations between pairs of items that are presented during learning are guaranteed to be adjacent items in the order, and observed that learning of an order can be reduced in this case to solving a linear regression task. Achieving through learning a representation of ranks in neural networks with a basically 1-dimensional dynamics, like that of area LIP, has been addressed besides (Di Antonio et al., 2024) only with the multilayer neural network model of (Nelli et al., 2023), where the activation of neurons on a hidden layer exhibited a 1D dynamics after Backprop training.

We present a minimal model for learning to extract and represent ranks in area LIP that is not only consistent with a large body of experimental data, but is also supported by a rigorous mathematical theory. The simplest possible assumption about the underlying network architecture is to have direct synaptic connections between neural populations that represent items and a neural system that has an LIP-like 1-dimensional dynamics as in (Ganguli et al., 2008), or even a single linear neuron. We show that this simple architecture suffices for learning rank estimates, with a simple and biologically plausible rule for synaptic plasticity. Furthermore, this learning capability is supported by a rigorous mathematical theory, the Rank Convergence Theorem. Our simple model exploits the commonly observed fact that different objects and concepts tend to represented in higher brain areas by approximately orthogonal neural codes. For example, it was found that different concepts or items are represented in the medial temporal lobe (MTL) of the human brain through the firing of almost disjoint assemblies of concept cells (Fried, 2022), i.e., by approximately orthogonal high-dimensional (high-D) neural codes. Approximate orthogonality of neural codes was also found in the hippocampus of the rodent (Sun et al., 2025). Orthogonality of item representations, or even one-hot encoding, has also been commonly assumed in other models for order learning, see e.g. (Nelli et al., 2023).

We then address experimental data which show how the brain integrates information from different learnt order relations into a cognitive map (Xiao et al., 2025), or into a larger order (Nelli et al., 2023). We are not aware of any model for the former, and only of a multi-layer perceptron model trained through stochastic gradient descent with different learning rates for each pair for the latter (Nelli et al., 2023). We show that our approach provides simple and transparent models for both of these order integration tasks.

Apart from modeling how the brain learns and represents order relations, it is also important to understand how such learning processes can be implemented in energy-efficient neuromorphic hardware with simple on-chip learning rules. Whereas previous models based on training multi-layer networks by backprop were not suitable for that, our simple model is suitable for implementation through on-chip learning with memristors that can assume a fair number of resistance values, such as the ones described in (Li et al., 2018). Furthermore, autonomous generation of cognitive maps from several learnt order relations as we show for the data of Xiao et al. (2025) from the human brain, appears to be useful for enabling navigation and abstract inference in robots.

In the last section of the paper we address another important result on neural representations of rank representations in the parietal cortex of the human brain. Although the primarily 1-dimensional dynamics of area LIP suggests that 2D projections of fMRI recordings of rank representations should lie on a line, they are consistently curved. The question why this is the case was explicitly raised in (Nelli et al., 2023). We provide a rigorous explanation of this effect based on the fact that ranks are not only represented in a summation code, but also by heterogeneous rank selective neurons. We show that this representation entails that 2D projections of neural codes for ranks lie on a curve, rather than on a line. The same theory also explains other questions about curved 2D projections of high-D neural codes that were addressed in (Okazawa et al., 2021).

## 2 Results

### 2.1 Architecture of a simple model for learning relations in area LIP

We want to understand how neurons in area LIP can learn to produce rank estimates for items **a**_*i*_ in an order “*≺*” that are each encoded by some high-D input vector. We assume that information about this order “*≺*” is provided in a realistic piecemeal fashion, through unstructured sequences of order relations **a**_*i*_*≺* **a**_*j*_ for arbitrary pairs of items (that are not necessarily adjacent in the order). Producing rank estimates for items in this order is a non-trivial computing and learning task, because the rank of an item **a**_*i*_ is a latent variable that depends in a complex way not only on order relations that involve this item **a**_*i*_, but also on order relations for all other pairs of items.

This learning process can best be modeled as online learning process, instead of a batch learning process that is more common in machine learning, but also in models for learning order relations. In online learning, weight updates are only carried out upon errors or surprises, i.e., when information about a relationship **a**_*i*_ *≺* **a**_*j*_ disagrees with current internal rank estimates, see Fig. 1A. The number of these errors is a standard measure for the efficiency of an online learning algorithm. Our model can also be trained using batch learning, and the standard measure for the efficiency of that, the number of examples that are presented presented—yields roughly twice the number of weight updates observed in online learning, see section B.2 of the Supplement.

**Fig. 1:**
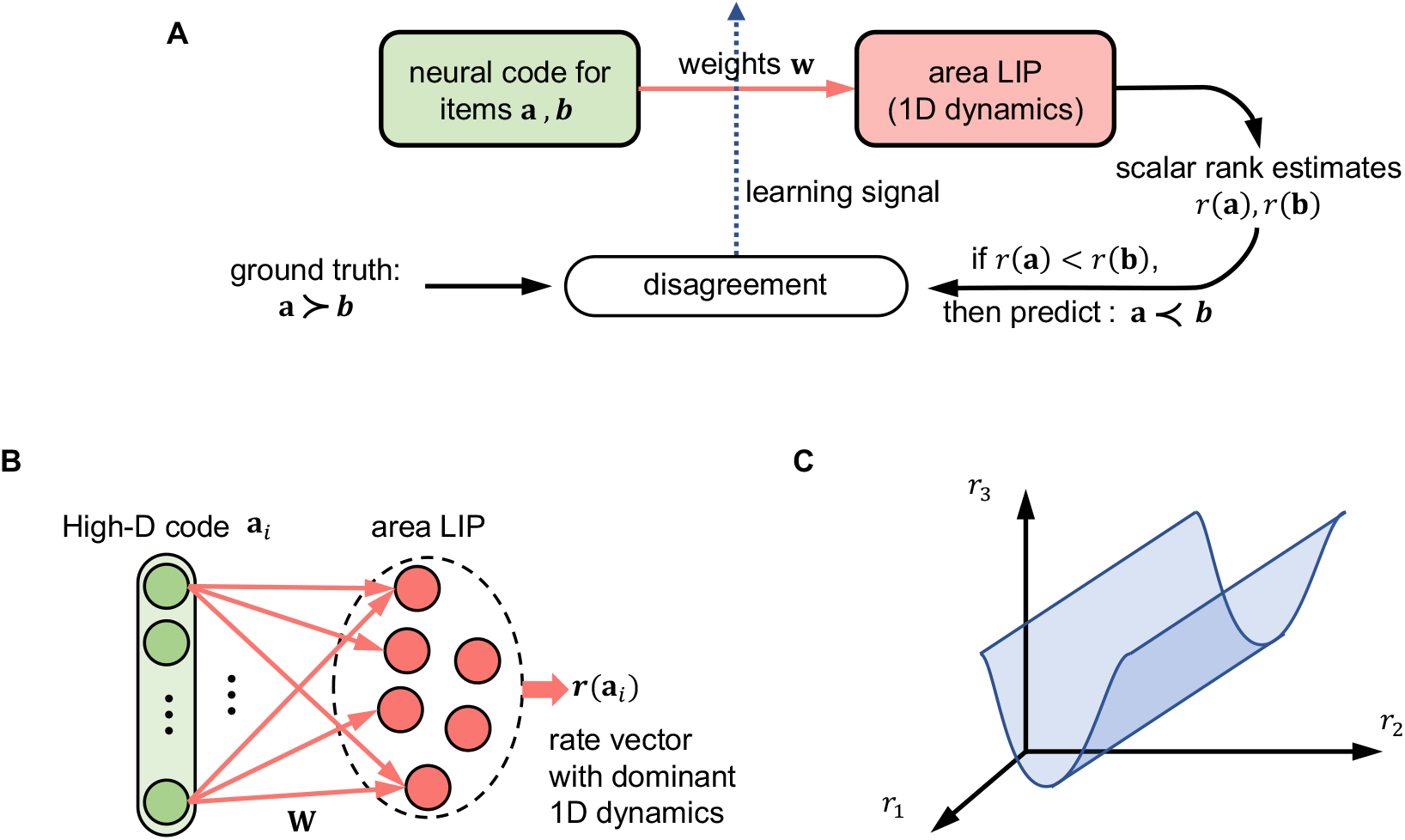
Architecture of our model. **A**. The underlying online learning framework: Weights are updated whenever the model produces through its rank estimates *r*(**a**) and *r*(**b**) predictions about the order between two items **a** and **b** that turn out to be wrong. **B**. Proposed architecture for learning rank estimates (summation code) in area LIP: High-D representations of items are projected through synapses with weights **w** onto a population of neurons that has an approximately 1-D dynamics. **C**. Scheme for 1-D dynamics of firing rates in a network: Activity states of the network drawn to a 1-D attractor a suggested by (Ganguli et al., 2008) (compare with their Fig. 2).

We show that this online learning process can be accomplished by a single layer of synapses with weights **w** to area LIP, as indicated in Fig. 1B. In other words, we show that after learning the overall firing activity **r**(**a**_*i*_) (space rate code) in a population of neurons that is targeted by these synaptic connections is able to provide consistent approximations of the ranks of all item **a**_*i*_ in a learned linear order “*≺*”.

According to the experimental data that we reviewed in the Introduction, the neural dynamics in area LIP is dominated by a 1-D line attractor (see Fig. 1C), that can be reproduced in a model through lateral connections between excitatory neurons (Ganguli et al., 2008). In order to make the learning process analytically tractable, we replace this population with 1-D dynamics as shown in Fig. 1B by a single linear neuron *n*, see Fig. 2A, that provides rank estimates for items **a**_*i*_ through its output *r*(**a**_*i*_). One can interpret it as firing rate of neuron *n* relative to a non-zero baseline firing rate, in order to account for negative values that it may assume.

**Fig. 2:**
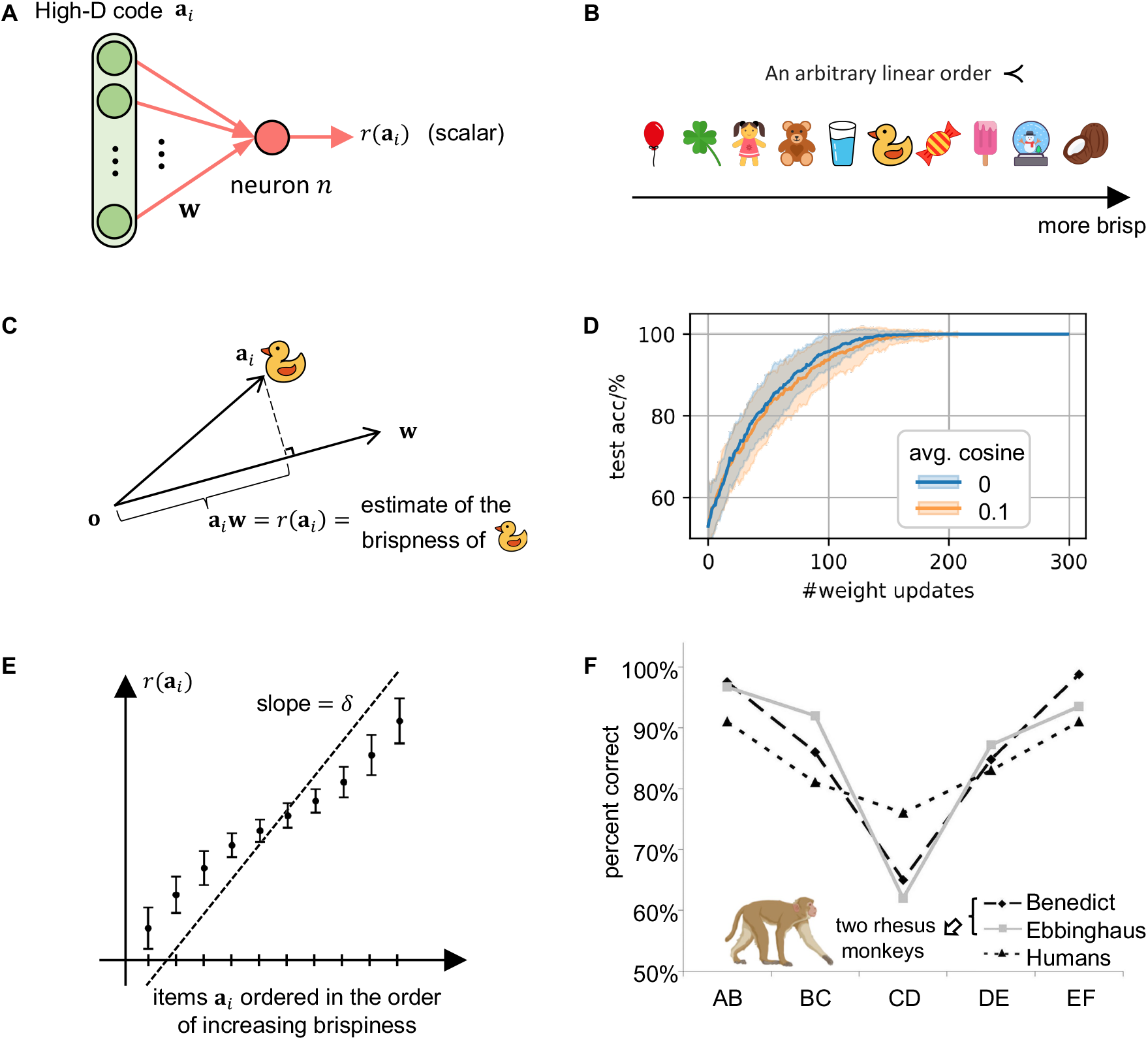
Performance of the base model, and comparison with brain data. **A**. Analytically tractable version of the model in panel B, with a single linear neuron representing the firing activity of a population with 1-dimensional dynamics. **B**. Task illustration: Learning the relation “more brisp than” for 10 items. **C**. The brispiness prediction for an item **a**_*i*_ (here: a duck) by the linear neuron of panel D can be understood geometrically as the length of the projection of the high-D neural code for **a**_*i*_ onto the weight vector **w**. Hence learning of **w** amounts to finding a direction of the weight vector where the lengths *r*(**a**_*i*_) of all projections of neural codes for items **a**_*i*_ onto **w** is consistent with their brispiness relation. **D**. Emergence of transitive inference capability. The accuracy of the model was evaluated after each weight update on non-adjacent pairs of items, although the learner had only received information about adjacent pairs. Perfect orthogonality of item representations **a**_*i*_ was assumed for the blue curve, and approximate orthogonality, suggested by experimental data, with an average cosine of 0.1 for the orange curve. One sees that the model does not require perfect orthogonality for fast learning. Means and standard deviations were calculated for 100 rounds of training using different random seeds. **E**. Brispiness predictions for all items were evaluated at that stage in the learning process when they became accurate (that took on average 93.6 weight updates). The dots and error bars indicate the means and standard deviations, respectively, computed over 100 rounds of training using different random seeds. The resulting rank estimates have a characteristic S-shape. **F**. Experimental data from (Merritt and Terrace, 2011) for two monkeys (“Benedict” and “Ebbinghaus”) and human subjects on the terminal item effect. Correctness of order estimates for pairs was much higher for pairs involving items at the end of the order, as predicted by the S-shape in panel E for the firing rates of neurons after learning. This panel is copied from Fig. 2 of (Merritt and Terrace, 2011).

**Fig. 3:**
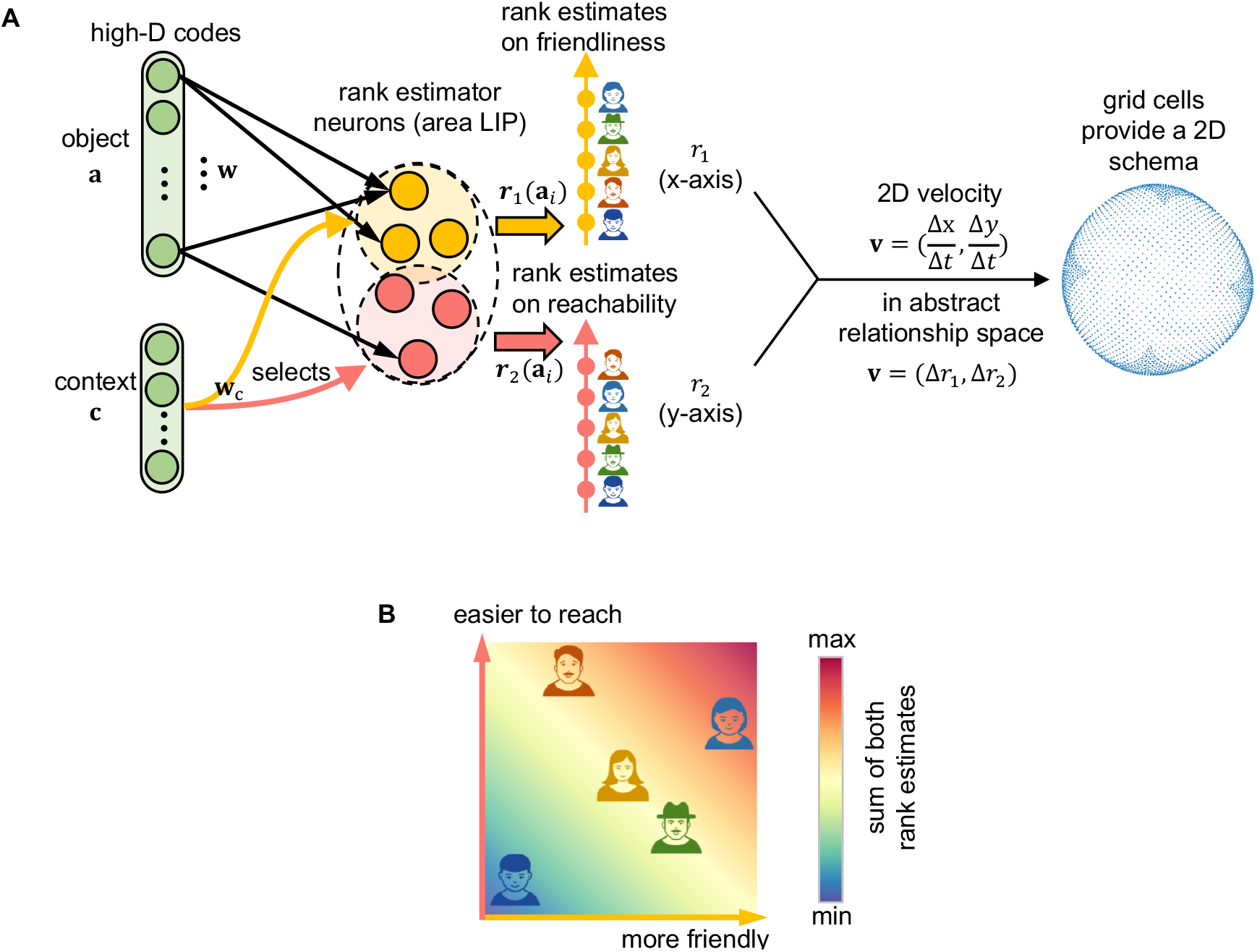
Emergence of a cognitive map from two learnt order relations. **A**. Rank estimation is computed as the sum of inputs from object coding (weights **w**) and from context coding (weights **w**_*c*_). The context code can then select subgroups of neurons to respond to a specific context by inhibiting the others. This yields parallel representations of ranks in two different orders by different neurons. One can treat the outputs of these neurons like velocity signals from area EC in a navigation task (Sorscher et al., 2023), and feed them into a grid cell network that creates from them a 2D cognitive map. Possible responses of grid cells can be plotted as locations on a torus (Gardner et al., 2022). The 2D schema shown here is the PCA result of grid cells’ activation for 50 50 mesh-grid locations in a small square that corresponds to an approximately flat fragment of the surface of the torus. **B**. An example of a 2D cognitive map that enables inference about five people in two relations (friendliness and reachability).

We apply the following simple learning rule for the weights of this neuron, to which we refer in the following as Rank Learning Rule:

#### Rank Learning Rule

If information is received that **a**_*i*_ *≺* **a**_*j*_ for some items **a**_*i*_ and **a**_*j*_, but **wa**_*j*_ is not by some given margin *δ* larger than **wa**_*i*_, one updates **w** by adding a fraction *η* of **a**_*j*_ and subtracting a fraction *η* of **a**_*i*_:

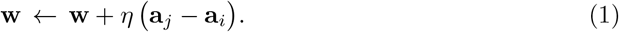

The parameters *δ* and *η* have fixed positive values, representing the margin and learning rate, respectively. In our experiment, the default values are *δ* = 1 and *η* = 0.1.

This learning rule is arguably the simplest and most straightforward one because it arises from gradient descent for the ranks of two fixed items **a**_*i*_ and **a**_*j*_, see section A of the Supplement. Hence, if it is sufficiently often iterated, it will eventually achieve that *r*(**a**_*i*_) + *δ < r*(**a**_*j*_). But updates for this pair of items are in general interwoven with updates for other pairs, which may contain some of the same items or not. Hence it is not a-priori clear whether this simple learning rule converges, i.e., arrives at rank estimates that are consistent with all order information that has been received. For example, the weight updates might get caught in infinite loops, since an item **a**_*i*_ has to satisfy many relations **a**_*i*_ *≺* **a**_*k*_ and **a**_*l*_ *≺* **a**_*i*_ with other items **a**_*k*_, **a**_*l*_. As long as the rank of **a**_*i*_ is not yet properly reflected in the output value of *n* for synaptic input **a**_*i*_, the Rank Learning Rule may sometimes add a fraction of **a**_*i*_ to **w** and sometimes subtract a fraction of it. These updates could potentially cancel each other. We show in Fig. 2B-C that the Rank Learning Rule works very well for learning the nonsensical relation “more brisp than” for 10 items (borrowed from (Nelli et al., 2023). For example, the item “candy” is more brisp than the item “yellow duck”. The neuron learns from a stream of information that one item is more brisp than some other item, and updates its internal model for predicting brispiness when needed, as indicated in Fig. 1A. The learning performance is shown in Fig. 2D. After, on average, 93.6 updates it produces rank estimates as shown in Fig. 2E that are consistent with the given order.

Transitive inference can be carried out by downstream networks in a straightforward manner on the basis of rank estimates that are encoded by firing rates in area LIP: In order to determine whichof two items comes later in the order they just have to evaluate which of them elicits a higher firing rate. This model is consistent with the commonly observed “symbolic distance effect” (Moyer and Bayer, 1976) in the performance of transitive inference by animals and human subjects: The accuracy of transitive inference increases, and the response time decreases, with the distance of two items in a learned order. Our model suggests that this effect arises because the comparison of two firing rates by downstream networks takes less time and is more reliable if the two firing rates are further apart.

The concrete rank estimates that emerge through the Rank Learning Rule, shown in Fig. 2E, provide an explanation for another commonly observed effect in transitive inference by human subjects: The “terminal effect” or “serial position effect” (Merritt and Terrace, 2011; Brunamonti et al., 2016; Mannella and Pezzulo, 2024). This effect describes the fact that transitive inference for pairs of items in a learned order has higher accuracy and requires less response time, compared with other pairs of items that have the same rank distance, if one of the two items is an endpoint in the learned order. For illustration, experimental data from (Merritt and Terrace, 2011) are shown in Fig. 2F). The subjects had learnt order relations between 6 images, labeled A, …, F through presentations of image pairs with adjacent labels, and were subsequently tested on image pairs with adjacent labels. One clearly sees better performance for pairs at the two ends of the linear order, This effect can be explained by the S-shape of the firing rates that emerge through learning with the Rank Learning Rule according to Fig. 2E: Firing rates for items at the endpoints of the learned order relation are further apart from those for next-ranked items, compared with distances between firing rates for other pairs of items that have the same rank difference. This non-linearity arises from the fact that the rank estimates for items at the endpoints are only pushed towards the outside by the Rank Learning Rule, whereas items in the mid-range are subject to numerous conflicting updates that push their rank estimates up and down.

The Rank Learning Rule works best if the items **a**_*i*_ are encoded by orthogonal vectors. Orthogonal codes have recently been found to arise commonly in the rodent hippocampus (Sun et al., 2025) But experimental data suggest that visual objects and concepts are represented also in the human brain by approximately orthogonal neural codes: Through sparse assemblies of “concept cells” in the MTL (Fried, 2022). If one encodes these assemblies by binary vectors, where a “1” indicates that a neuron belongs to this assembly, these experimental data support a model where items are encoded by sparse high-D binary vectors. These vectors can not be assumed to be perfectly orthogonal, since assemblies for different concepts are known to have an overlap between 1% and 4%, depending on the degree by which the underlying concepts are associated (De Falco et al., 2016). Fig. S1A shows that this amount of overlap between assemblies amounts to a cosine similarity between 0.02 and 0.08 for the corresponding binary vectors. The orange curves in Fig. 2D, Fig. 4D-E, and Fig. S1B show that the learning performance is still very good for an average pairwise cosine value of 0.1 between vectors that represent different items. In other words, the level of orthogonality that is needed for the Rank Learning Rule is reached by the approximate orthogonality that has been found in neural codes for concepts in the MTL of the human brain, no matter whether they are associated or not.

**Fig. 4:**
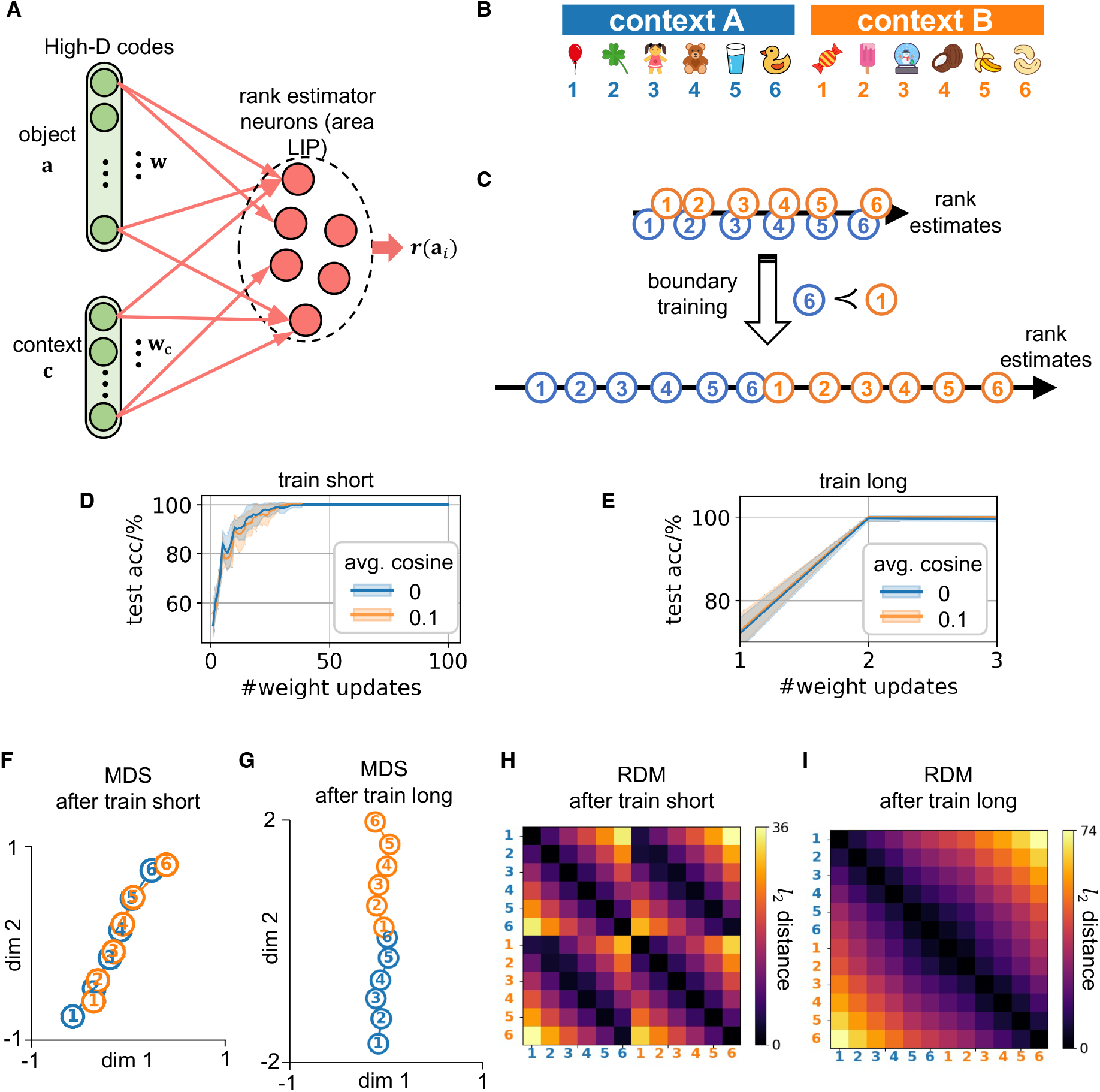
Combining information from separately learned orders. **A**. Network architecture: Rank estimation is given by the sum of inputs from object coding (weights **w**) and from context cod- ing (weights **w**_*c*_). **B**. Task illustration: Two linear orders are learned separately in different contexts A and B during “train short”. **C**. In a 2nd learning phase “train long” the information is provided that the item with lowest rank in context B comes after the item with the highest rank in context A (“boundary pair”). Resulting rank estimates of our model after train short and train long are shown. **D**. Learning curve of the 1^st^ learning phase “train short”. Model accuracy was evaluated after each weight update on all item pairs within the same context, while the learner received information only about the adjacent pairs within each context. **E**. Learning curve of the 2^nd^ learning phase “train long”. Model accuracy was evaluated after each weight update on all item pairs in the combined order, while the learner received information only about the boundary pair. Two presentations of the boundary pair were sufficient. The blue curves in both **D** and **E** assume perfect orthogonality between item representations **a** and context representations **c**, while the orange curves assume approximate orthog- onality with an average cosine of 0.1. Mean and standard deviations in both **D** and **E** were computed from 100 training sessions with different random seeds across train-short and train-long conditions. The model learns rapidly even with just approximate orthogonality, reaching 100% accuracy in all sessions after just two updates. **F**. and **G**. 2D projections via MDS (Multidimensional scaling) of the response of the rank estimator neurons from panel A for the items from the two contexts after train short and after train long, respectively. **H**. and **I**. Representational dissimilarity matrices (RDMs) capture pairwise rank estimation distances as *RDM*_*i,j*_ = *∥***r**(**a**_*i*_) *−***r**(**a**_*j*_) *∥*_2_ after short and long train- ing, respectively. The same neural representation as for panels **F** and **G** was used.

We are not aware of experimental data which elucidate whether a transitive inference query “is **a**_*i*_ *≺* **a**_*j*_?” is processed by neural circuits of the brain by estimating the ranks of these two items in a sequential manner, or whether these items are provided simultaneously as input to these neural circuits. Similarly, it is not clear whether order information **a**_*i*_ *≺* **a**_*j*_ is processed during learning by placing attention sequentially on each of the two items, or in a fully parallel manner. The sequential process can be supported by the argument that placing attention simultaneously on two different items appears to be difficult, at least without explicit training. Our model is neutral with regard to this issue, since it can also be used for fully parallel computations: By giving neural codes for both items simultaneously as inputs to the linear neuron, but **a**_*j*_ with factor +1 and **a**_*i*_ with factor *−*1, as in the models of (Nelli et al., 2023; Di Antonio et al., 2024). After learning, the linear neuron will then output the rank difference of both items, and this value is positive if and only if the answer to the question “is **a**_*i*_ *≺* **a**_*j*_?” is “yes”. Note that also the Rank Learning Rule is compatible with this fully parallel model, and the weight update is in this case a fraction of the total input **a**_*j*_ *−* **a**_*i*_ that it receives.

Fig. 2D-E show that learning rank estimates for brispiness works very well in our simple model. But it is not clear whether this example is representative. A deeper analysis of the dynamics of the learning process is needed in order to determine whether it converges for any given order, and for arbitrary sequences of pieces of information that inform about the order relations between some pairs of items. This analysis is provided in the following section.

### 2.2 Learning theory for the Rank Learning Rule

A simple geometrical argument shows that there exists for any order and any encoding of items by orthogonal vectors a weight vector **w** which enables the neuron *n* to produce accurate rank estimates for all *N* items in the order. Fig. 2C suggests that it suffices to make sure that each item has a projection of a suitable length onto the vector **w**. Hence one can simply define it for any correct set of ranks *r*_*i*_ for the items **a**_*i*_ as

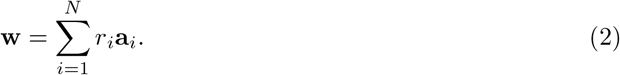

But this simple arguments, assumes that one has already correct rank estimates *r*_*i*_, although these are not provided directly during the learning process. Furthermore, it does not entail that the Rank Learning Rule converges to a vector **w** that provides correct rank estimates. It had been shown in (Di Antonio et al., 2024) that for the special case where only pairs of items are presented during learning that are adjacent in the order that is to be learnt, learning of the order can be reduced to a linear regression task. The optimal solution can be approximated via gradient descent, where the gradient descent rule is identical to the Rank Learning Rule, except that it requires a varying continuous-valued factor that measures continuous-valued deviations from target values. In contrast, we show that the Rank Learning Rule reaches after finitely many steps an optimal solution with a fixed learning rate. Importantly, it learns an order also if non-adjacent pairs of items are presented during learning. This can not be achieved if order learning is formulated as regression task, because one would then have to provide the actual rank difference of the two items, which becomes only known after the order has been learnt.

The learning theory for the Rank Learning Rule is governed by a rigorous mathematical result, the Rank Convergence Theorem. This Theorem guarantees that the Rank Learning Rule converges for every given total order “*≺*”, and for any order of presentations of pairs of items during learning, after finitely many weight updates. The precise statement of the Rank Convergence Theorem (see Methods) is analogous to that of the famous Perceptron Convergence Theorem, which guarantees convergence of online learning with a simple plasticity rule (the delta learning rule) for learning binary classifications. But the Rank Learning Rule has to solve a more difficult task, since it does not receive target ranks by a teacher, which would correspond to the target classifications which the learner receives in the case of the Perceptron Convergence Theorem. Rather, it has to learn a latent variable, the rank, which has no analogue when one learns classifications. Consequently, one cannot adopt the proof the the Perceptron Convergence Theorem in order to prove the Rank Convergence Theorem. Rather, one needs a new argument for the proof. We present this argument in full detail in the Methods section.

Note that the Rank Convergence Theorem only guarantees that if one keeps on presenting pairs of items for which the estimated ranks are not yet consistent with their order relation, this process will end after finitely many steps with a weight vector **w** for which the rank estimates that neuron *n* provides are fully consistent with the order. But if one stops this learning process prematurely, the rank estimates that neuron *n* provides are in general not yet consistent with all order relations.

There may even be cases where the order relations between some pairs of items are unknown, e.g. when a partial order is to be learned. In this case the Rank Convergence Theorem guarantees that the learning process converges to rank estimates that are consistent with all pairwise order relations that exist in the partial order. In order to apply the Rank Convergence Theorem, which assumes existence of a total order that is consistent with all examples, one can use the fact that every partial order can be embedded into a total order.

An empirical finding is that the inference of new relations between items—those implied by other previously revealed relationships—reaches high accuracy quite fast (see Fig. 2D and Fig. 4D-E). The y-axes of these plots illustrate the emergent correctness of these inferences during learning. Furthermore, as detailed in Sec. 2.4, our model learns approximately 7–24× faster than humans in one experiment and 10× faster in another one under identical settings. This may be due to the presence of noise in neural circuits of the human brain, or to rules for synaptic plasticity that require more iterations. But one can adjust our learning model to these behavioral data by lowering its learning rate.

### 2.3 Emergence of cognitive maps from several learnt order relationships

In the real world, an item may participate in multiple order relations. For example, we learn that some people are more friendly than others, and independently, that also some of them are easier to reach than others, see Fig. 3A for an illustration. The recent study (Xiao et al., 2025) shows through neural recordings from the human brain that it integrates information from two learnt orders into a 2D cognitive maps. Compared with transitive inference for a single order, this cognitive map enables a much larger repertoire of abstract inference tasks that involve both orders. We show that a transparent model for these experimental data arises if one duplicates our base model for learning two linear orders so that the firing rates of two different neurons (or two populations of neurons) in area LIP provide rank estimates for an item in the two orders. Integration of information from two different contexts in area LIP had already been examined in (Kumano et al., 2016). We capture that by letting context inputs gate off irrelevant neurons via strong inhibition through **w**_*c*_, see Fig. 3A.

Routing the outputs of neurons that report rank estimates for the two linear orders to a standard model for a grid cell system from (Sorscher et al., 2023) yields a 2D map that supports navigation and complex inference by combining information from both orders. The vector of the two rank estimates plays here a role that is analogous to the 2D velocity vector that provides for 2D navigation input to grid cells in the entorhinal cortex (EC) (Sorscher et al., 2023).

To see how the resulting cognitive map enables inference that combines information from both order relations, we consider in Fig. 3B the query “Who is easiest to reach and most friendly?”. Within our model, this reduces to identifying the item closest to the top right corner of the cognitive map. But the geometry of this cognitive map also supports completely different inference tasks, such as finding the person that has the largest mismatch between friendliness and reachability.

### 2.4 Rapid reconfiguration of the internal model when separately learned orders are combined on the basis of new information

Humans can rapidly reassemble knowledge about separately learned orders, for example merge two separately learned orders on the basis of new information. This process was studied in experiments with human subjects in (Nelli et al., 2023). They first learned two separate orders in phase 1, and in phase 2 that the first item in one order comes after the last item in the other order, i.e., the learned orders were appended to each other, see Fig. 4B-C.

The Rank Convergence Theorem guarantees that the Rank Learning Rule will also lead in this 2-phase learning process to rank estimates that are in the end consistent with all order relations for the combined order. We show that if area LIP receives besides synaptic input that encodes items also synaptic input that encodes the contexts A and B to which the items of the two short orders belong (see Fig. 3A and Fig. 4A), then fast reconfiguration of the internal model emerges (Fig. 4E). It is well-known that higher brain areas, especially the PFC, keep track of the current context, and signal it via spike inputs to virtually all other brain areas (Heald et al., 2023). Furthermore, it is pointed out in that review that context is typically encoded in the PFC by firing patterns of neurons that are orthogonal to neural codes for other task-relevant variables (Bernardi et al., 2020). This was recently confirmed for the human brain in (Courellis et al., 2023).

In order to model rapid configuration of internal we complemented the Rank Learning Rule by a similarly simple Context Learning Rule for the weights of synaptic connections that project context information to area LIP (see Fig. 4A):

#### Context Learning Rule

If information is received that **a**_*i*_ *≺* **a**_*j*_ for some items **a**_*i*_ and **a**_*j*_ that come from two different contexts A and B and are encoded by neural codes **c**_*A*_ and **c**_*B*_, but (**wa**_*j*_ + **w**_*c*_**c**_*B*_) is not by some given margin *δ* larger than (**wa**_*i*_ + **w**_*c*_**c**_*A*_), one also updates the weights **w**_*c*_ for neural codes for context by adding a fraction *η*_*c*_ of **c**_*B*_ and subtracting a fraction *η*_*c*_ of **c**_*A*_:

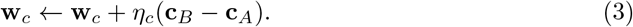

We used random vectors **c**_*A*_ and **c**_*B*_ to encode contexts and applied a larger learning rate *η*_*c*_ = 8*×η* to the context weights **w**_*c*_ compared to the other weights **w**. Like *η*, the value of *η*_*c*_ is kept constant throughout the training process. We tested the resulting learning model for synaptic connections to area LIP on the scenario from (Nelli et al., 2023). First, two different orders were learned in contexts A and B, see Fig. 4B, using the same weight vectors **w** and **w**_*c*_ for learning both orders. The learning curve is shown in 4D. For train long, we continued learning for these weight vectors with a boundary pair, consisting of the item that came last in the short order from context A and the first item in the other short order from context B. One sees in Fig. 4E that in general 2 presentations of the boundary pair suffice for adjusting all rank estimates for the long order.

MDS projections of the representations of the items after train short and after train long are shown in Fig. 4F-G. Hence this simple model leads to very similar results as the model of (Nelli et al., 2023) for the same experiment, although the architecture and learning processes were very different in the two models. (Nelli et al., 2023) employed a multi-layer neural network that was trained through backpropagation (BP). Their model required in addition that on the side also a certainty matrix was learned for all pairs of items, that was not part of the neural network representation. It was needed to set differential learning rates for all pairs of items. Neither BP nor certainty estimates are needed in our model. In addition, the reconfiguration of ranks for the long order was substantially faster: two presentations of the boundary pair were sufficient, see Fig. 4E.

### 2.5 Explaining curved 2D projections of neural representations of linear orders in the human brain

One structural difference between the neural network representation of a learned order in the brain and in the multilayer perceptron (MLP) model of (Nelli et al., 2023) had been addressed in the Discussion section of that study: 2D projections of fMRI data from the human brain have a characteristic horseshoe shape, i.e., they are curved (inflected). In contrast, 2D projections of the MLP representations of items in a learned total order from (Nelli et al., 2023) tend to form a line. Fig. 4F-G show that this also occurs in our model. However, we have considered so far only 2D projections of responses of neurons that encode ranks by a summation code. The brain encodes ranks also by neurons that fire most strongly for particular ranks or regions of ranks (heterogeneous code) (Summerfield et al., 2020). We show here that the projection of their activity into 2D is necessarily horseshoe shaped, more precisely, the projected points are predicted to approximate a parabola, provided the neurons have some overlap in their preferred ranks.

A transformation of the summation code that emerges from the previously discussed learning rules into a heterogeneous code is straightforward and requires no new learning. It suffices that the outputs of summation coding neurons provide synaptic inputs to an array of excitatory neurons with staggered firing thresholds. Each of these neurons will fire then only for ranks above a certain value. If these neurons are interconnected with inhibitory neurons so that several negative feedback loops arise (each enclosed by dashed lines in Fig. S3), very high ranks will activate this negative feedback so strongly that the neurons no longer respond to very high ranks. As a result, each excitatory neuron in the array become tuned to some interval of ranks. The width of this tuning is regulated by the strength of inhibition: According to (Nieder, 2016) a blockage of GABAergic inhibition leads to a broadening of the tuning profiles of these excitatory neurons. Hence, a network architecture as indicated in Fig. 5A, see Fig. S3 for a more detailed plot, leads to a diversity of tuning profiles of soft rank selective neurons with different approximately Gaussian tuning profiles, as indicated in the middle of Fig. 5A. The matrix on the right side of that panel records in each column the rank values for which one of these neurons is selective, in terms of a firing rate being above some given threshold. Essential for our theoretical analysis is that these Gaussian tuning curves overlap, in this example by two rank values. Therefore we refer to them as soft rank selective neurons. The key point from the mathematical perspective is that the covariance matrix that is generated by these soft rank selective neurons can be approximated by a circulant matrix, i.e., by a matrix where successive rows are shifted by one column, see Methods or (Gray et al., 2006) for a precise definition.

**Fig. 5:**
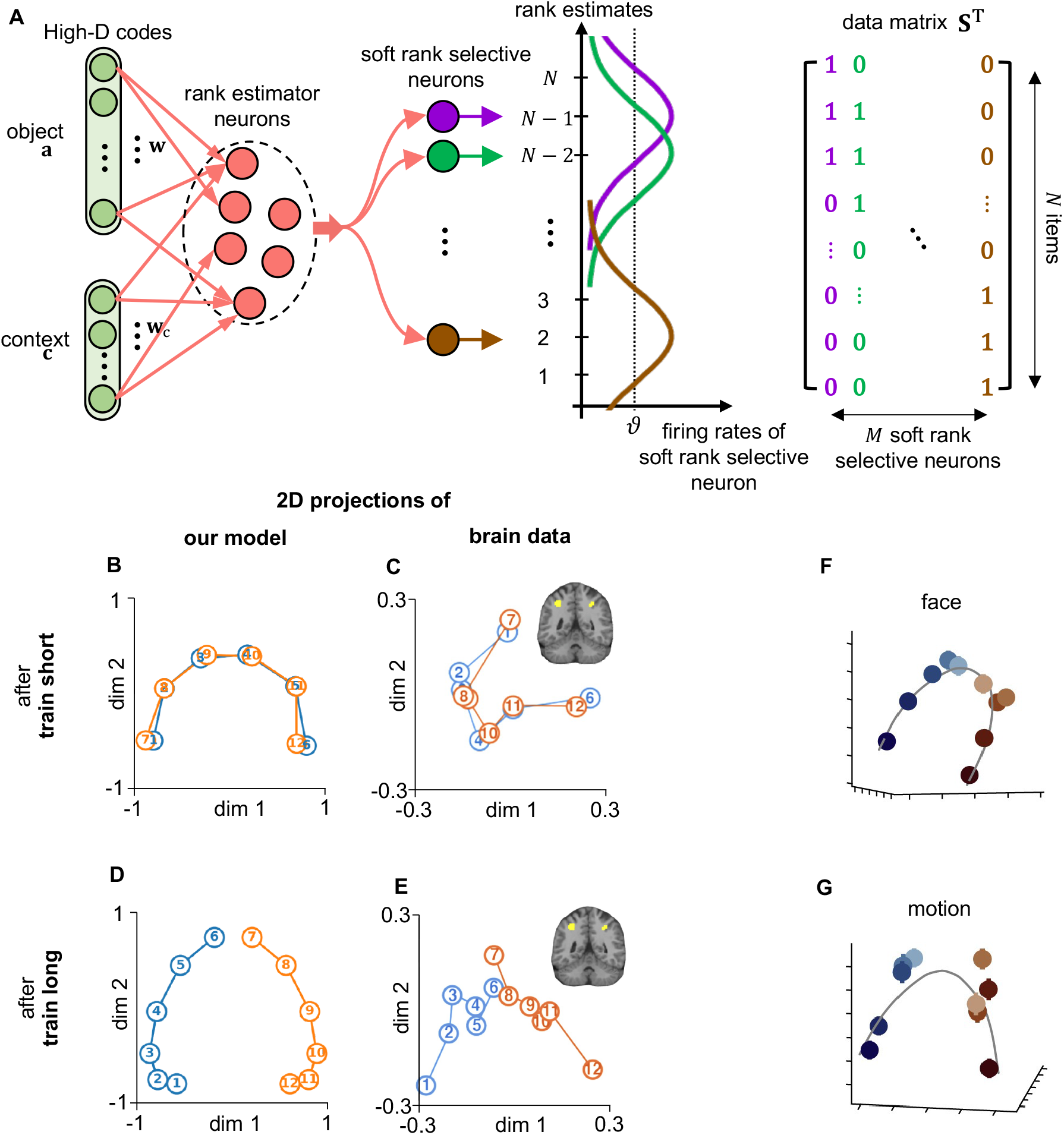
Emergence of a parabola (horseshoe) shape of 2D PCA (principal component analysis) projections through soft rank selective neurons. **A**. Architecture for the generation of heterogeneous codes from a summation codes for rank estimates. Each colored circle in the middle indicates a population of excitatory neurons with negative feedback from inhibitory neurons, see Fig. S3 for a more detailed plot. This negative feedback curtails the firing rate in each population. On the other hand, staggered firing thresholds of excitatory neurons in different populations cause different onsets of their preferred ranks. Hence, the response profile of each population can be approximated by a Gaussian. If one binarizes their firing responses via some threshold, one arrives at the simple binary matrix on the right, which we analyze in Section 4.5 of Methods and Section E of the Supplement. **B**. Shown are projections onto the first two PCA dimensions, i.e., onto the first two eigenvectors of the covariance matrix, of the patterns of firing activity of soft rank selective neurons for 6 items in the two separately learnt short orders. This is shown here after *training short*, but before *training long*. **C**. Corresponding 2D projection of human BOLD signals of area PPC for the same order (Copied from Fig. 3C of (Nelli et al., 2023)). **D**. 2D PCA of the same neural data after *training long*, revealing a characteristic horseshoe. **E**. Matching BOLD projection post-*training long* ((Nelli et al., 2023) Fig. 5B). In (Nelli et al., 2023), the 2D projection method is multi-dimentional scaling (MDS), which usually generates similar results as PCA in this case. **F-G**. PCA projections of multi-unit activity in area LIP of the monkey for two linearly ordered sets of stimuli, “face” and “motion”, 400 ms after stimulus onset (copied from Fig. 2 E-F of (Okazawa et al., 2021)). The gray lines are cubic smoothing splines fit. Error bars are standard error of the mean.

Our theoretical analysis of the reasons for horseshoe shaped 2D projections of neural activity is based on the structure of eigenvectors of circulant matrices: They can be approximated by the cosine function, see section 4.5 of Methods. Of particular interest for the analysis of the first two dimensions of a PCA projection are the projections of neural codes onto the first two eigenvectors, for which these approximations via the cosine function are shown in Fig. 6. More precisely, projections onto the first eigenvector can be approximated by the cosine function over 180 degree. Projections onto the second eigenvector can be approximated by the cosine function over 360 degrees. Using the double- angle formula, cos(2*θ*) = 2 cos^2^(*θ*) *−* 1, one can approximate the shape of the 2D PCA projection by a parabolic function. This yields the horseshoe-shaped inflection in Fig. 5B and D. These theoretically predicted 2D projections provide a qualitative fit to the recorded data from the human brain, as reported in (Nelli et al., 2023), see Fig. 5C and E. In Section 4.5 of Methods and Section E of the Supplement we provide a rigorous proof of this effect. Note that the signs of eigenvectors can be chosen arbitrarily. Hence PCA projections of the same data that are flipped upside down or left to right are equally valid.

**Fig. 6:**
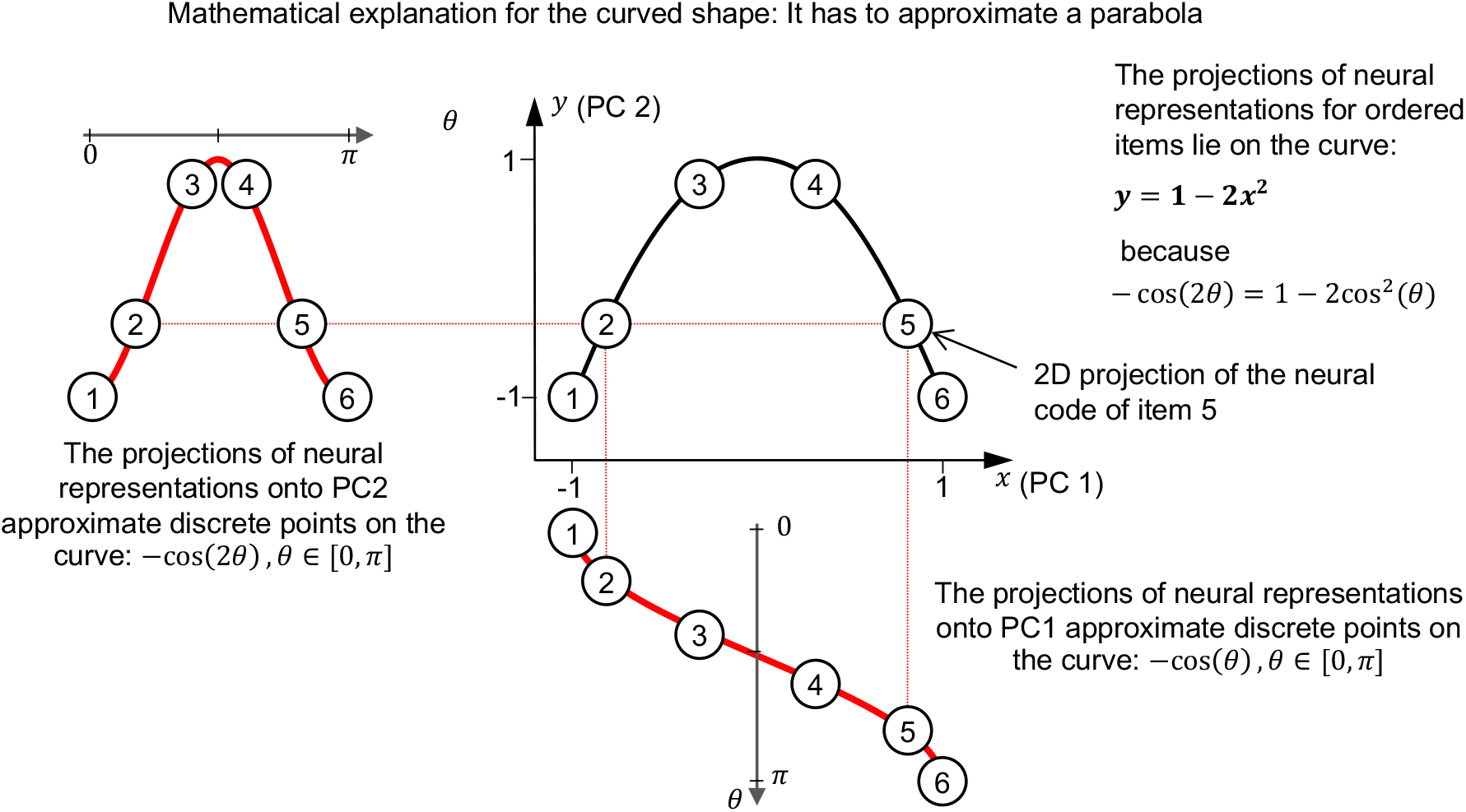
Mathematical illustration of the cause of the curved shape. According to Eq. (33) and (Eq. (34)), the projections of the firing activity of soft rank selective neurons onto the first principal component (PC1) approximate the cosine function over [0, *π*], see the lower part of the plot, while the projections on PC2 approximate the cosine over [0, 2*π*], see the plot at the left side. Combined, they come to lie on a parabola. This parabola explains why the horseshoe shape emerges in the PCA results of fMRI data. Note that we flipped the sign of both eigenvectors so that the PCA result resemble those in (Nelli et al., 2023).

A horse-shoe shape of 2D projections was also found for multi-unit electrode recordings from area LIP of the macaque for linearly ordered items (Okazawa et al., 2021), see Fig. 5F and G. Their emergence can also be explained by our theory. These panels also show that the approximation of the covariance matrix by a circulant matrix is already good enough for this small instance (with *N* = 6 items and *M* = 8 soft rank selective neurons) so that its eigenvectors of the covariance matrix behave qualitatively like those of a circulant matrix.

## 3 Discussion

We have shown that the all-important latent variable for learning an order, the rank of an item in this order, can be learned by a simple plasticity rule for synaptic connections from neurons that encode items or context to neurons in area LIP that participate in the commonly observed representation of learned ranks through a summation code (Summerfield et al., 2020; Bottini and Doeller, 2020), i.e., by a space rate code (see Fig. 1A). This latent variable has to be extracted from dispersed information about order relations between different pairs of items, and it is not a-priori clear whether this is possible. But the convergence of this extraction process is guaranteed by a new theoretical foundation, the Rank Convergence Theorem. The same online learning method may also explain the emergence of rank estimates in other brain areas such as the PFC (Brunamonti et al., 2016).

The emergent S-shape of resulting rank estimates in our model (Fig. 2E) provides a principled explanation for the commonly observed “terminal effect” or “serial position effect” in human behavior, whereby transitive inference performance is enhanced when the comparison involves items located at the end of a linear order, see (Merritt and Terrace, 2011; Brunamonti et al., 2016; Mannella and Pezzulo, 2024). Using rank estimates for transitive inference also explains the commonly observed “symbolic distance effect”, whereby transitive inference performance increases linearly with the distance of items in the learned order. This effect arises automatically when transitive inference is based on rank estimates that are encoded in an internal model by firing rates: If the firing rates that encode the rank estimates for two items have a larger distance, this difference can be detected faster and more reliably by downstream networks.

If one takes into account that LIP neurons also receive information about the context of the current relational learning process, one can explain by our model also the emergence of a cognitive map that enables navigation in a 2D space that is spanned by ranks in two unrelated order relations, as shown for the human brain in (Xiao et al., 2025). Our model can also explain rapid reconfiguration of rank estimates in the light of new order relations, as addressed in the experiments and model of (Nelli et al., 2023). An interesting difference between the learning process in their multi-layer perceptron model and our simple model is that our model does not require Backprop training, and also no differential regulation of learning rates for different pairs of items is required for reconfiguration of learned order representations. As a result, substantially fewer learning steps are needed by our model for the reconfiguration of learned order relations in the face of new information.

Our model for learning order relations gives rise to several experimentally testable predictions. One prediction concerns the connectivity: It predicts that approximately orthogonal neural representations of order items in other brain areas are projected by synaptic connections to LIP neurons, whose firing rates encode after learning estimates for ranks through a summation code. Our model also predicts that these LIP neurons that signal rank estimates through their firing rate provide the primary input to downstream networks that carry out transitive inference. This would explain the terminal item effect (Mannella and Pezzulo, 2024) on the basis of inherent biases of their firing rates (see Fig. 2E). Furthermore, our model predicts that the weights of synaptic connections to these LIP neurons undergo synaptic plasticity during learning of an order. It also predicts that weight updates primarily occur after an error or surprise, i.e., when a presented order relation is not predicted by the emergent internal model. Hence, similar as for BTSP (Behavioral Time Scale Synaptic Plasticity) and other plasticity rules that have been demonstrated in-vivo in the adult brain (Magee and Grienberger, 2020; Chéreau et al., 2022), 3rd factors that signal surprise are predicted to play a more important role than postsynaptic activity for learning order relations by these LIP neurons. In addition, our model predicts that information about the behavioral context of the currently learned order is also transmitted by synaptic connections to the same LIP neurons, and that the weights of these synapses are subject to similar rules for synaptic plasticity. Our model for the experimental data of (Xiao et al., 2025) predicts that outputs of LIP neurons that learn rank estimates are routed to the hippocampal formation, and have a similar impact on grid cell firing as velocity vectors in the case of locomotion.

In the last part of the paper we have proposed a theoretically founded solution to an open problem that was addressed in (Nelli et al., 2023): 2D projections of internal representations of ranks in the parietal cortex of the human brain are not straight, as predicted by most models, but on a horseshoe shaped curve. We have shown that if one takes into account that ranks are not only represented by summation codes of neurons, but also by soft rank selective neurons that preferentially fire for a certain range of ranks, the experimentally observed inflection of 2D projections automatically emerges. It results from the structure of the first two eigenvectors of circulant matrices that arise in the mathematical analysis of neurons with soft selectivity for ranks. In fact, this analysis is applicable also to neurons with soft selectivity for some other magnitude, thereby also providing a new and principled explanation for the experimentally observed curved manifolds for decision variables in the macaque brain (Okazawa et al., 2021). Our theory predicts that horseshoe shaped curvature will arise in all 2D projections of brain activity that represents value of some scalar variable, provided that there exist neurons with soft selectivity for values of this variable.

The Rank Learning Rule that emerged from our normative approach is consistent with experimental data on synaptic plasticity in the brain, especially with more recent data from in-vivo experiments, see the reviews (Magee and Grienberger, 2020) and (Chéreau et al., 2022). It has been shown there that synaptic plasticity in the living and behaving animal depends primarily on presynaptic firing and on a gating signal, and less on the timing of postsynaptic spikes as proposed by the STDP (Spike-timing-dependent plasticity) rule (Markram et al., 1997). More specifically, the Rank Learning Rule that we have examined requires LTP (long-term potentiation) in order to increase the rank estimate for a synaptic input **a**_*j*_ and LTD (long-term depression) for decreasing the rank estimate for some synaptic input **a**_*i*_. It had been shown in (Hong et al., 2022) that separate eligibility traces for LTP and LTD exist, and that their impact on weight updates can be selectively gated by neuromodulatory signals.

Altogether, we have provided a new theoretical foundation for understanding how order relations are learnt and represented in the brain, and how these representations can give rise to intelligent decision making. These processes appear to be instrumental for enabling our brains to cope with a web of relations that impinge on us both from the environment and in the form of learn relations between abstract concepts. Our theory suggests that the computational and learning processes which are required for that are facilitated by inherent capabilities of neurons and synaptic plasticity that we have elucidated in this work. The stability and convergence of these learning processes is guaranteed by mathematical facts.

Apart from understanding brain processes, our theoretical foundation also provides algorithmic and architectural guidance for building energy-efficient edge devices that are able to cope with a web of relationships that impinge on them. It shows that fast learning and representations of multiple types of relations is possible that are easy to combine, for example in cognitive maps, in a way that supports low-latency decision making. Furthermore, the Rank Learning Rule that we have presented is within reach of on-chip learning through memristor arrays, such as for example the ones described in (Li et al., 2018).

## 4 Methods

### 4.1 Details to the Rank Learning Rule in Section 2.1

Here we introduce a loss function whose gradient descent optimization exactly reproduces the Rank Learning Rule.

Consider a fixed pair of items represented by coding vectors **a**_*i*_ and **a**_*j*_, where **a**_*i*_ *≺* **a**_*j*_, with a margin *δ >* 0 defining the minimal desired separation between their rank estimates. Let the firing rates (rank estimates) of the rank estimator neuron for these items be defined as *r*(**a**_*i*_) = **wa**_*i*_ and *r*(**a**_*j*_) = **wa**_*j*_.

We define a pairwise ranking loss function as follows:

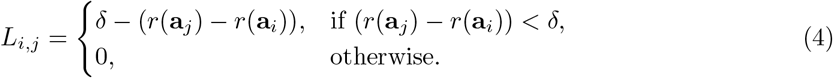

Minimizing this loss function (See eq. (4) in Supplementary Sec. A. for detail) via gradient descent leads precisely to the weight update given by the Rank Learning Rule:

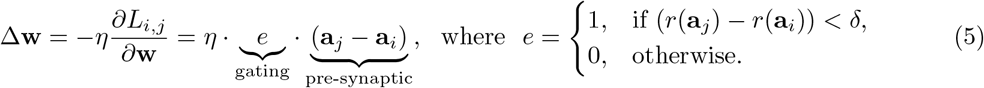

### 4.2 Proof of the Rank Convergence Theorem and details of experiments, discussed in Section 2.2

#### Rank Convergence Theorem

Let **a**_1_, …, **a**_*K*_ be some arbitrary finite set of non-zero orthogonal vectors, to which we refer as items. We consider updates of weights **w** under the Rank Learning Rule:

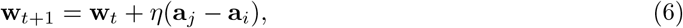

for arbitrarily selected pairs of items **a**_*j*_ and **a**_*i*_ with the property that **a**_*i*_ *≺* **a**_*j*_ but **w**_*t*_**a**_*j*_ is not by some margin *δ* larger than **w**_*t*_**a**_*i*_. The factor *η* is assumed to be some arbitrary positive fixed value.

Then **w**_*t*_ converges to some finite value, independently of the order and frequency by which pairs of items **a**_*j*_ and **a**_*i*_ are selected for updates.

**Proof:**

Assume, for contradiction, that the claim is false, i.e., that the learning process does *not* terminate, and infinitely many updates are performed by the Rank Learning Rule.

We consider the subset of items

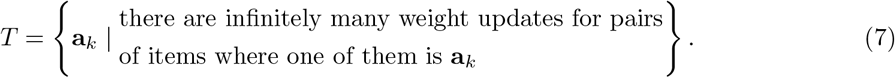

By assumption, *T* is nonempty. We pick an element **a**_min_ *∈ T* that is minimal under the ordering *≺*.

**Observation 1: w**_*t*_**a**_min_ *→ −∞*.

Since only finitely many updates are made for items **a**_*k*_ *≺* **a**_min_, from some step *t* on, all weight updates that involve **a**_min_ also involve some **a**_min_ *≺* **a**_*k*_. Hence from some step *t* on only negative updates are made for **a**_min_, i.e., *η***a**_min_ is subtracted from **w** whenever a weight update is made that involves **a**_min_. Hence, the orthogonality of the items **a**_*i*_ implies that **w**_*t*_**a**_min_ *→ −∞*.

**Observation 2:** A dual version of this argument shows that

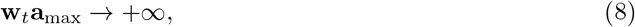

for the maximal item **a**_max_ in *T* with regard to *≺*.

**Observation 3:** Let item **a**_*k*_ in *T* have the property that **w**_*t*_**a**_*k*_ *→ −∞* and **w**_*t*_**a**_*l*_ *→ −∞* for all **a**_*l*_ in *T* with **a**_*l*_ *≺* **a**_*k*_. If **a**_*q*_ is the next larger item in *T* with regard to *≺*, then also **w**_*t*_**a**_*q*_ *→ −∞*.

This holds because after a while, all updates of **w** where *η***a**_*q*_ is added to **w**_*t*_ are made at steps *t* when **w**_*t*_**a**_*q*_ *≤* **w**_*t*_**a**_*r*_ + *δ* for some **a**_*r*_ in *T* with **a**_*r*_ *≺* **a**_*q*_. Since by assumption **w**_*t*_**a**_*r*_ *→ −∞* for all these **a**_*r*_, the products **w**_*t*_**a**_*q*_ that occur at these steps also converge to *−∞*.

Induction along *≺*, where the induction beginning is provided by Observation 1 and the induction step by Observation 3, implies then that **w**_*t*_**a**_*k*_ *→ −∞* for all **a**_*k*_ in *T*. But this yields a contradiction to Observation 2.

This completes the proof of the Rank Convergence Theorem.

#### 4.2.1 Details to learning of a given order in Fig. 2

In Fig. 2B, we trained our model on adjacent pairs among the 10 items, and tested on unseen non-adjacent pairs among the same set of items. The learning rate *η* in Eq. (5) is set to 0.1. Each item code **a**_*i*_ is a 10^4^ dimensional random binary vector with number of ones uniformly sampled within the range [0.2%, 1%]. The blue curve is measured when all **a**_*i*_ forms an orthogonal set; and the orange curve is measured when the average cosine similarity between all pairs of items’ code is 0.1. Each synaptic weight in **w** is initialized by a normal distribution *𝒩* (0, 10^*−*4^). The weights are initialized very small to prevent unnecessary random biases that decreases the learning speed. The standard deviations of the two curves are calculated by repeated training from scratch 100 times. The randomness comes from data sampling, as a random pair of adjacent objects is picked at each training step. The margin *δ* is set to 1.

#### 4.2.2 Method for generating codes for items with a given average pairwise cosine similarity

After initializing each item code with a fixed number of ones, this number is kept constant for each item. The indices are adjusted either to overlap with the dimensions of other items’ ones, increasing the average cosine similarity, or to shift overlapping ones to different dimensions, reducing the cosine similarity. For a given target average cosine similarity, this process is iterated for a sufficiently long time to achieve a numerically optimal coding configuration.

### 4.3 Details to our model for the emergence of cognitive maps from several learnt order relationships, discussed in Section 2.3

#### 4.3.1 Context selection

The architecture in Fig. 3 requires distinct subgroups of rank-estimator neurons to be selected for encoding item ranks under different contexts, a property reported by (Kumano et al., 2016). This can be implemented in multiple ways. Here we show arguably the simplest method: Using fixed, high-dimensional sparse-binary context codes together with a randomly initialized high-D sparse-binary weight matrix **w**_*c*_, each context is automatically routed to a distinct, non-overlapping subgroup of LIP neurons without any synaptic learning. Other learning rules that introduce competitive mechanisms between LIP neurons—such as lateral inhibition or winner-take-all (WTA)—may also achieve the same effect.

Let each context *c ∈ {* 1, …, *K}* be represented by a binary vector 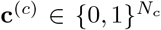, and assume approximate orthogonality:

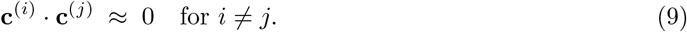

For a randomly initialized sparse-binary weight matrix 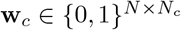 (sparsity *p ≪* 1, mapping input vectors of dimension *N*_*c*_ to *N* LIP neurons) whose *N*_*c*_ columns satisfy that any two distinct columns are also approximately orthogonal:

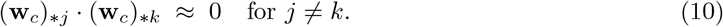

Upon presentation of context **c**, the net drive to the downstream population is

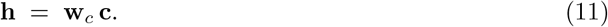

One sees that **h** is exactly the sum of the columns of **w**_*c*_ selected by the non-zero index of **c**.

Because the context codes are orthogonal and the columns of **w**_*c*_ are orthogonal,

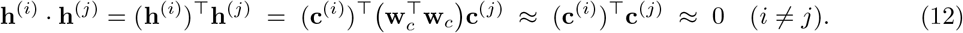

Therefore, each context selects approximately distinct subgroup of neurons. When these context inputs induce subthreshold depolarization in specific neuronal ensembles, subsequent object-driven signals can then selectively recruit different LIP subpopulations based on that context.

#### 4.3.2 PCA of grid cells

The grid cells naturally form a representation of any 2D space. In Fig. 3 A, we showed a 2D projection of grid activations on a (4 *×* 4) square environment via PCA.

The grid cells are randomly initialized. With 1000 grid cells, each initialized independently and modeled as the sum of three planar cosine gratings 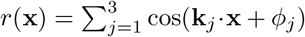, whose wave-vectors **k**_*j*_ are 60^*°*^ apart and share magnitude |**k**_*j*_| = 2*π/λ*. The grid scale was drawn once per cell from *λ ∼ U* (0.05, 8). To eliminate directional bias a global rotation *θ*_0_ *∼ U* (0, 2*π*) was applied, giving arg(**k**_*j*_) = *θ*_0_ + *jπ/*3.

The PCA is performed on the grid activations of 50 *×* 50 evenly sampled mesh-grid locations on the square environment.

### 4.4 Details to combining separately learned orders into one order, discussed in Section 2.4

#### 4.4.1 Architecture

When generate the MDS plot in Fig. 4F-G, we use a population of rank estimator neurons **n**’s activations to encode rank information collectively. The population in this experiment contains 100 rank estimator neurons. These rank estimator neurons receive summation inputs from both codes, with each neuron’s value *r*:

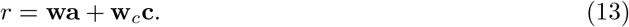

Each neuron learns independently its incoming synaptic weights based on the same learning rule Eq. (5) as a single rank estimator neuron. The symmetry breaks because of the different initialization with each synaptic weights sampled from *N* (0, 10^*−*4^).

#### 4.4.2 Train short

During initial training (“train short”), we divided the 12 objects into two distinct sets (context A 1-6 and context B 1-6) and presented in alternating trails. The model only made comparisons within each context and indicate which of two adjacent-ranked objects (e.g., context A 3 and context A 4) was larger or smaller, receiving fully informative feedback. In total, the model is trained on 80 pairs of examples to form the results in Fig. 4F and H. This training allowed our model to infer ranks within each set (i.e., context A 1–6 and context B 1–6) but did not reveal any information about the rank relations between the two sets (e.g., context A 2 *<* context B 3). We set the learning rate *η* = 0.1. Given that training occurs only between items from the same context, no weight updates to **w**_*c*_ would occur based on the context learning rule.

#### 4.4.3 Train long

In (Nelli et al., 2023), human participants underwent 20 training trials (each contains a pair of object together with their ranking information) after being informed that the lowest ranked object “6” in context A is ranked lower than the lowest ranked object “1” in context B. We replicated this setting, and reproduced their results by training our model on this boundary pair for 2 times, only 1*/*10 of the total training trials of the human participants. The result is shown in Fig. 4G and I.

The key of assembling two order is to shift the rank estimation of a whole group together, without altering the in-context ranking, which, in our model, corresponds to prioritize the learning to happen on the context information than the item code. By analyze the contribution by different variables on the rank estimation changes after applying the weight updating in Eq. (5), with an assumption that codes are orthogonal, one gets:

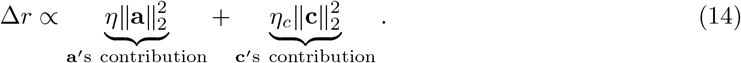

This result from our model suggests that two potential factors may contribute to the context- prior effect: 1. The learning rate for the context-related weights *η*_*c*_ is higher than the learning rate *η* for the object weights. 2. The *l*_2_-norm of the context code **c** is larger than that of the object code **a**. A detailed analysis of how these two factors influence the prioritization of learning based on context information is provided in the supplementary information Sec. C. In our experiment, we set *η*_*c*_ = 0.4 and *η* = 0.05; the dimension *D* = 10^4^ of **a** equals to the dimension of **c**; the sparsity of **a** equals to the sparsity of **c** with 0.2% to 1% (sampled from the uniform distribution) neurons fire in each code, which is based on the data from human medial temporal lobe (Fried, 2022; Waydo et al., 2006).

### 4.5 Theoretical analysis of the shape of 2D projections of the firing activity of soft rank selective neurons for items in an order, discussed in Section 2.5

We analyze the application of PCA for reducing the dimensionality of the representation of *N* items by *M* soft rank selective neurons from *M* to 2. The goal of this section is to derive a closed-form solution for the projection results.

The structure of this section follows the steps of computing PCA:

1. Calculation of the covariance matrix
2. Approximation and eigenvector calculation of the covariance matrix
3. Calculation of the projections of neural representations onto the first two principal components

#### 4.5.1 Calculation of the covariance matrix

Assume that we have *M* soft rank selective neurons *s*_1_ to *s*_*M*_ for items **a**_1_ to **a**_*N*_ in an order. We denote in the following the response of neuron *s*_*i*_ to item **a**_*j*_ as **s**_*i*_(*j*).

We now consider the case where

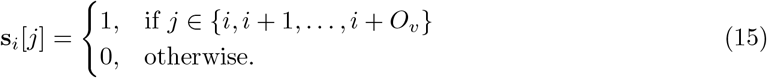

These are the columns in the example matrix **S** as shown in Fig. 5A. In this equation, *O*_*v*_ (overlap) represents the number of items for which a soft rank-selective neuron fires beyond its central rank.

The responses of all *M* soft rank selective neurons to the *N* items is encoded by the matrix

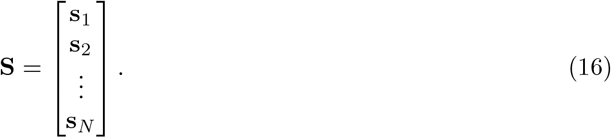

Firstly, we compute the mean vector of the responses of the soft rank selective neurons to different items:

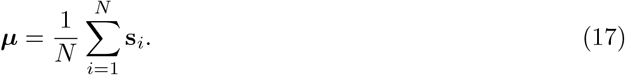

We then compute the centered data matrix:

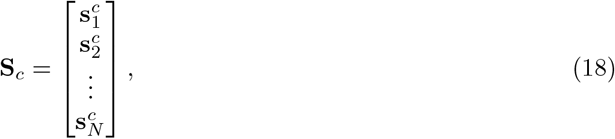

where each response vector is centered by subtracting the mean vector:

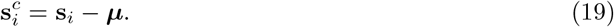

The covariance matrix **C** *∈* ℝ*M ×M* is defined by:

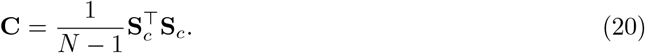

The variance terms in the main diagonal of **C** are:

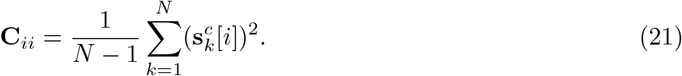

The covariance terms are:

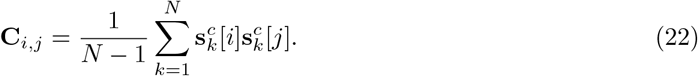

The principal components are the first two eigenvectors of **C**. However, without simplification, it is difficult to obtain a closed-form solution. Hence, we will introduce some assumptions to simplify the equations in the following sections.

#### 4.5.2 Approximation and eigenvector calculation of the covariance matrix

We show that **C** is a banded symmetric Toeplitz matrix, and hence can be approximated by a circulant matrix. Then we use properties of the eigenvectors of circulant matrices.

##### Assumption

Assume *N* is reasonably large so the edge effect can be neglected. Specifically, we assume that:

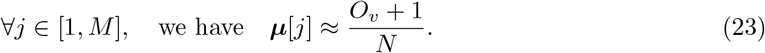

This equation holds rigorously for items that are not at an end of an order.

The centered data vectors then can be approximated by:

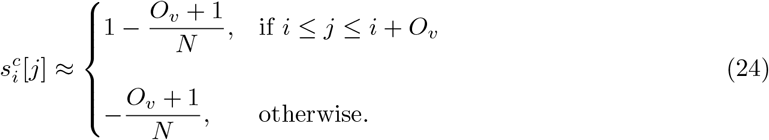

Since 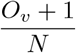 is small for large *N*, we can further approximate:

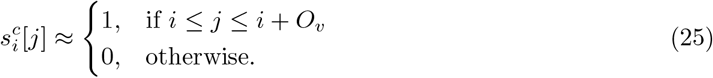

Under our assumption, the covariance matrix **C** is symmetric and Toeplitz, meaning that **C**_*i,j*_ depends only on |*i− j* |.

The covariance between dimensions *i* and *j* is:

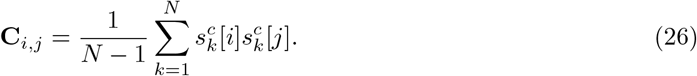

Given the approximated centered data, 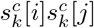 is non-zero only when neural representations of items overlap in dimensions *i* and *j*. Hence, we can approximate **C**_*i,j*_ by:

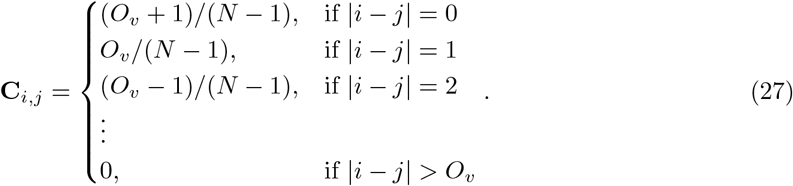

The form of the resulting covariance matrix is the **banded symmetric Toeplitz matrix** that satisfies the following properties:

- **Toeplitz structure:** The matrix has constant values along each diagonal, i.e., **C**_*i,j*_ = **C**_*i*+*k,j*+*k*_ for all *i, j, k* such that the indices are within bounds.
- **Symmetry:** The matrix is symmetric, meaning **C**_*i,j*_ = **C**_*j,i*_.
- **Banded structure:** The matrix has nonzero entries only within a fixed number of diagonals. Specifically, there exists a bandwidth *b ≥* 0 such that **C**_*i,j*_ = 0 whenever |*i− j*| *> b*. We have *b* = *O*_*v*_ in our setting as depict in Eq. (27).

An example **C** is provided below:

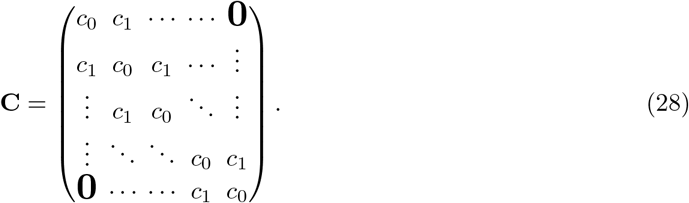

A banded symmetric Toeplitz covariance matrix can be approximated by a **circulant matrix C**^*′*^ *∈* ℝ*M ×M* (Gray et al., 2006), which, by definition, has a:

- **Cyclic row structure:** Each row of **C**^*′*^ is obtained by shifting the preceding row one element to the right (wrapping around at the boundaries). If the first row is

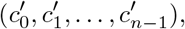

then the *i*-th row is given by

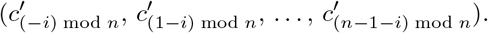

If one approximates the example banded symmetric Toeplitz matrix **C** above by a circulant matrix **C**^*′*^, then **C**^*′*^ would have the following form:

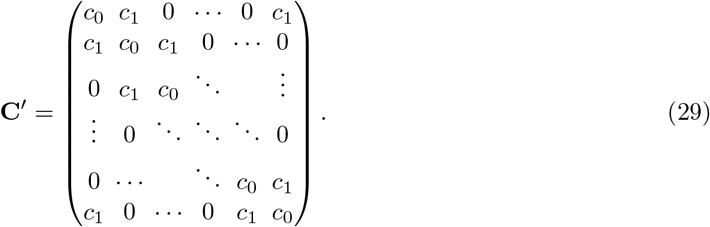

The only difference compared to **C** is that **C**^*′*^ requires an additional elements *c*_1_ in the upper right and lower left corner to fulfill the circulant property. Using a circulant matrix to approximate a banded Toeplitz matrix becomes more accurate as the size of the matrix *M* grows and the bandwidth *O*_*v*_ becomes narrower.

Importantly, a circulant matrix has **eigenvectors that can be represented by the columns of the discrete Fourier transform (DFT) matrix** (Gray et al., 2006). In the case of a real symmetric circulant matrix, the DFT eigenvectors can be re-expressed in a real-valued form. Specifically, the discrete cosine transform (DCT) provides an orthogonal basis that yields a closed-form representation of the eigenvectors (Britanak et al., 2010):

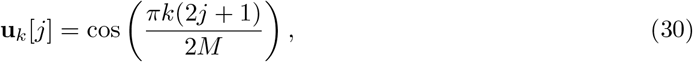

where *k* = 1, 2, …, *M* is the mode index, and *j* = 1, 2, …, *M* is the position index of each eigenvector. The proof of the validity of this step can be found in the supplementary information Sec. E.

We focus on the first two principal components, which correspond to the eigenvectors with the largest two eigenvalues (usually corresponding to *k* = 1 and *k* = 2):

- **First eigenvector**

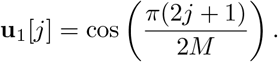

- **Second eigenvector**

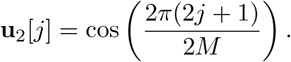

#### 4.5.3 Calculation of the projections of neural data onto the first two principal components

The projection of 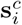 onto the *k*-th principal component is:

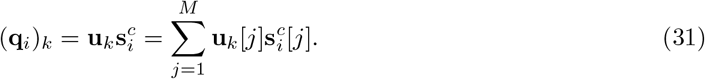

Substituting the approximated **u**_*k*_[*j*] and 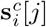 from eq. (30) and eq. (25), we get:

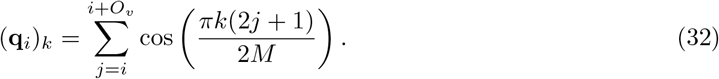

An intuitive observation is that the projection result can be viewed as a windowed sum of cosine values from *i* to *i* + *O*_*v*_, where *O*_*v*_ (overlap) represents the number of items for which a soft rank-selective neuron fires beyond its central rank. With *k* = 1 and under the assumption that the window is short (i.e., small *O*_*v*_), this windowed sum approximates a slightly smoothed cosine function over [0, *π*] for *i ∈* [0, *N*] (Fig. 6, *x*-axis). Therefore, one gets the projection onto PC1 as:

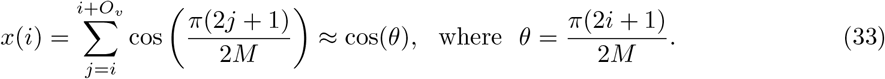

Similarly, the projection onto PC2 (*k* = 2) follows a discrete cosine approximation over [0, 2*π*] for *i ∈* [0, *N*] (Fig. 6, *y*-axis):

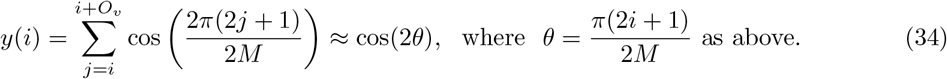

With (*x, y*) *≈* (cos(*θ*), cos(2*θ*)), where *θ ∈* [0, *π*], and using the trigonometric identity cos(2*θ*) = 2 cos^2^(*θ*) *−* 1, one can derive the explicit shape of the PCA results as:

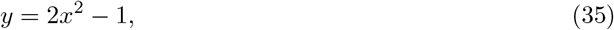

which follows a parabolic curve.

In addition, the property of eigenvectors allows the sign of each eigenvector to be chosen freely. To better align with the PCA results in (Nelli et al., 2023), we flipped the sign of the projection result on both axes: (*x, y*) *≈* (*−* cos(*θ*), *−* cos(2*θ*)) and get:

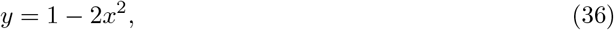

which explains the inflection effect in Fig. 6.

## Acknowledgments

We gratefully acknowledge the generous advice of Christopher Summerfield, and helpful comments by Giovanni Pezzulo, Guozhang Chen, Alice Dauphin, and Hui Lin on an earlier version of the ms. This research was partially supported by the National Science Foundation of the USA (EFRI BRAID project 2318152) and the Austrian Science Fund (FWF) (10.55776/COE12).

## Supplementary Information for

### A Rank Learning Rule as local gradient descent

We show that for a fixed pair of items **a**_*i*_ and **a**_*j*_ the Rank Learning Rule from Eq. (1) and Eq. (5) is equivalent to performing gradient descent on a corresponding loss function for these two items. This analysis suggests that the Rank Learning Rule makes sense, but makes no implication for the realistic scenario where many different pairs of items, some with overlap, are presented to the learner.

Given a pair of items **a**_*i*_ *≺* **a**_*j*_ with rank estimates *r*(**a**_*i*_) = **wa**_*i*_ and *r*(**a**_*j*_) = **wa**_*j*_, and a margin *δ >* 0, we define the loss function as:

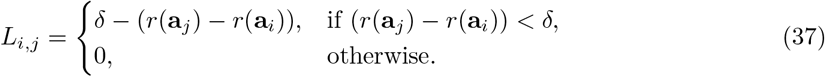

When calculating the gradient with respect to **w**, we obtain:

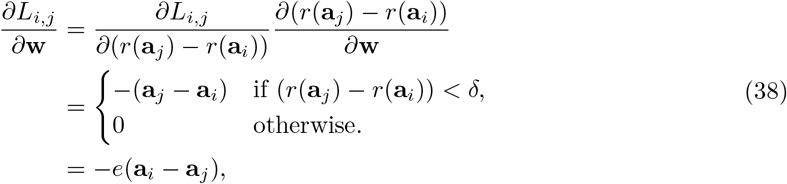

where

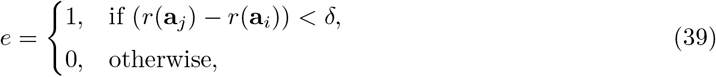

is the error signal.

Hence, applying gradient descent for a fixed pair of items results in an update rule that matches Eq. (5):

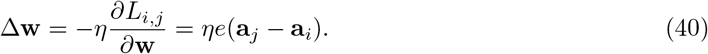

## B Further details to Sec 2.1

### B.1 Impact of relaxed orthogonality of neural codes for items

The brain represents items by assemblies of concept cells, that fire when this item is presented (or mentally considered) (De Falco et al., 2016). An important question for this study is to what extent these neural presentations of items can be viewed as being represented by orthogonal vectors. These assemblies are sparse (consisting of 0.2% to 1% of neurons in the human medial temporal lobe (Waydo et al., 2006; Fried, 2022)). But some neurons respond to more than one item. In other words, the assemblies have in general some overlap. The relative size of the overlap between two assemblies of concept cells for two different concepts has been estimated experimentally by the relative frequency that a neuron that responds to one concept also responds to the other concept. More precisely, the overlap between two assemblies is measured as % of neurons in their intersection relative to the size if the union of both assemblies.

**Fig. S1:**
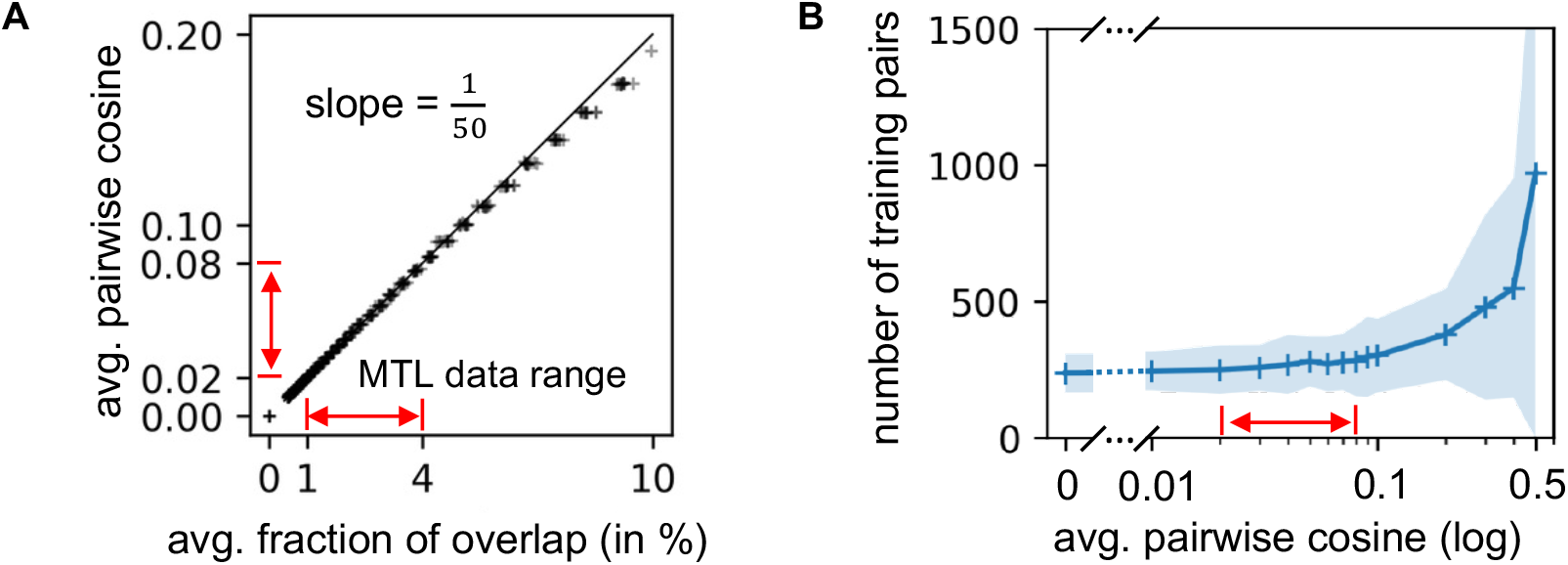
**A**. The average fraction of overlap and the average pairwise cosine similarity follow a linear relationship in the range of small overlaps that we consider. Data from the human medial temporal lobe (MTL) suggest that the average fraction of overlap between different objects falls within the range of 1% to 4% (De Falco et al., 2016; Fried, 2022). From our simulation results, we observe that this range corresponds to an average cosine similarity between 0.02 and 0.08. A linear regression fit shows that the average cosine similarity is approximately 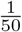 of the average fraction of overlap, as depicted by the black line. **B**. Number of pairs of items presented, required for the agent to reach 100% testing accuracy with varying average cosine similarity, for the task of learning brispiness among 10 items. Lines and shaded areas represent the averages and standard deviations from 100 repeated experiments. Training is relatively fast when the average pairwise cosine is *<* 0.1, aligning with human MTL data [0.02, 0.08]. (Note: The x-axis of **B** is marked with (log), it means the scale of the axis adopts the logarithmic scale. Values shown on its side are still the original values.)

We show in Fig. S1A empirically that the fraction of overlap between assemblies for two different items is (in the range of small overlaps) approximately linearly related to the cosine similarity between the two corresponding binary vectors that encode these assemblies. This empirical analysis was carried out by assigning a binary vector of dimension 10^4^ to each assembly (item), with sparseness uniformly sampled from the range of 0.2% to 1% of 1’s. In addition, Fig. S1A shows that overlaps between 1 and 4 % yield cosine values between 0.02 and 0.08. The detailed method used to adjust the average fraction of overlap is discussed in Sec. 4.2.2.

Subsequently, we show in Fig. S1B that the learning speed of our model is for the range of these cosine values between 0.02 and 0.08 almost the same as for perfect orthogonality, i.e. cosine value 0. The task is the same as in Fig. 2B, where a single total order is learned. All settings are the same as depicted in Sec. **??**.

### B.2 Relationship between the number of weight updates in an online learning model and the number of item pairs needed to achieve the same performance in batch training using the rank learning rule

We illustrated the standard online learning approach in Fig. 1A: weights are updated only when there is a “surprise”; otherwise, they remain unchanged. For this online learning rule, we use the number of weight updates (in Fig. 2D and Fig. 3D-E) throughout the paper to evaluate learning speed. However, this measure for online learning differs from the standard measure in batch learning, where one counts the number of training examples (each example consists of a pair of items being compared). In this section we examine empirically the relationship between these two different measures for learning speed for the “briskness” learning experiment.

**Fig. S2:**
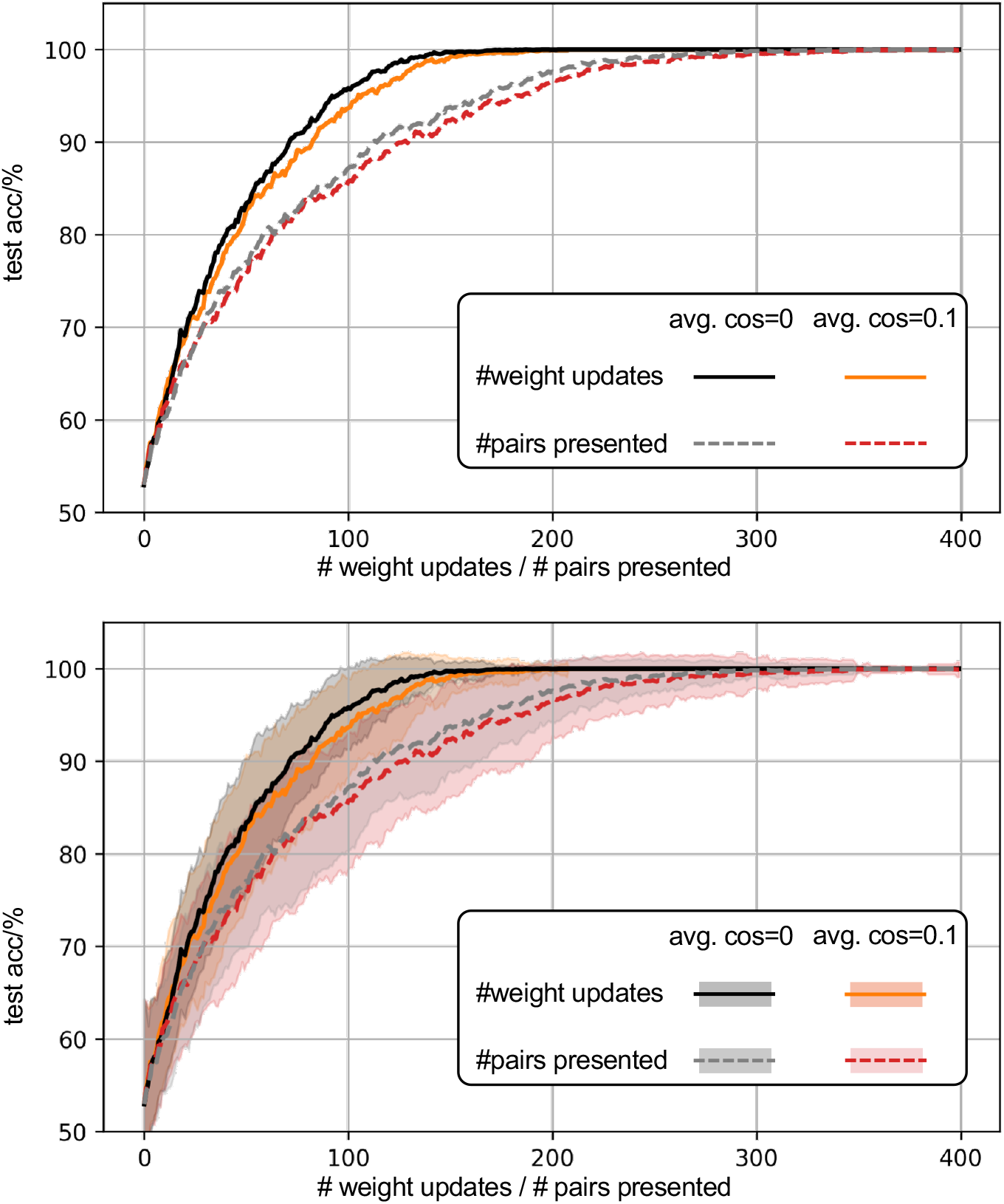
Relationship between the number of weight updates and the number of pairs presented (standard deviation omitted in the upper panel for better visual clarity). The pairs are accompanied by their relative ranking information, which indicates which item is ranked higher. The comparison is made under conditions of both perfect orthogonality and approximate orthogonality (with an average cosine similarity of 0.1).

From the figure above, it is clear that there is a difference between the two measures. During training from scratch until the results become accurate, only about 55% of the presented pairs trigger weight updates in online learning, meaning that roughly 45% of the pairs do not lead to any update, even when all pairs are presented with equal probability. This indicates that, throughout the training process until accuracy is achieved, weight updates occur for only slightly more than half of the training pairs.

## C Further details to Section 2.4

To understand how learning prioritizes the context code **c** over the item code **a**, we analyze how weight updates affect rank estimates.

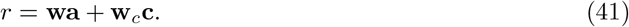

After updating the weights according to Eq. (5), and considering different learning rates, we have:

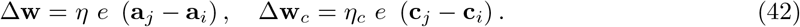

When the compared objects **a**_*i*_ and **a**_*j*_ are from the same context, the second term is zero, indicating that **w**_*c*_ does not participate in the “train short” phase.

The new rank estimate of objects **a**_*i*_ and **a**_*j*_ are:

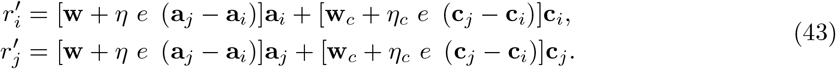

Because different brain codes are typically orthogonal to each other:

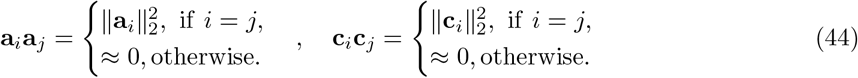

Therefore:

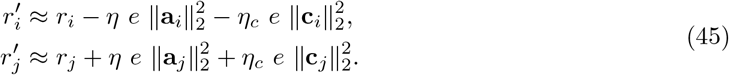

When calculating the rank estimate change, the factors affecting the rank estimate changes are:

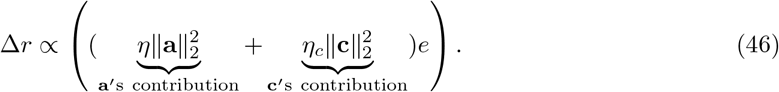

The above theoretical analysis clarifies the methods available for prioritizing learning in different representations. Whether learning happens more on the object embedding or context embedding depends on the ratio between:

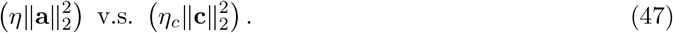

## D Neural network architecture that leads to the emergence of soft rank selective neurons, see Section 2.5

We show here the architecture that was indicated in Fig. 4A in more detail.

**Fig. S3:**
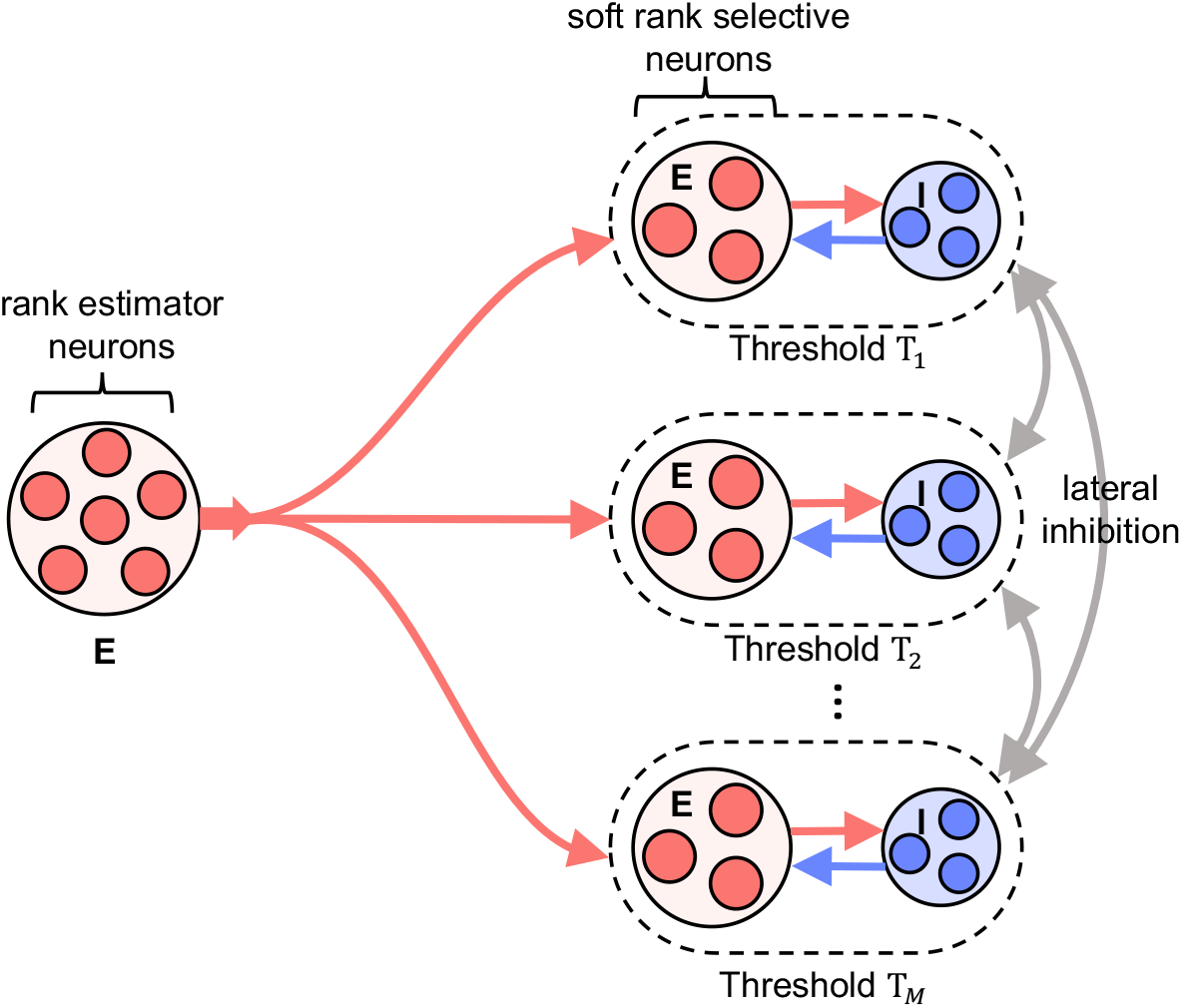
A population of rank estimator neurons (summation coding neurons) projects to various populations of other neurons. Each population consists of interconnected excitatory and inhibitory neurons, with distinct firing thresholds for excitatory neurons in different populations. Inputs below the threshold value for a population do not activate it. When the inputs exceed the threshold, excita- tory neurons are activated, but their firing activity is curtailed by negative feedback from inhibitory neurons. Inhibitory neurons may also provide lateral inhibition between populations in order to ensure that the overlaps in their tuning curves are limited.

## E Proof of Validity for the Eigenvector Approximations in Sec. 4.5

We need to show that:

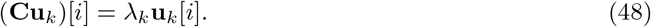

Using the Toeplitz structure:

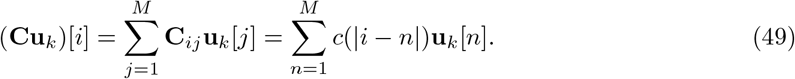

Substitute **u**_*k*_[*n*]:

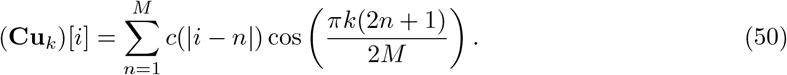

Let *m* = *n − i*. Then *n* = *i* + *m*, and *m* ranges from *−*(*i −* 1) to *M − i*. Substituting:

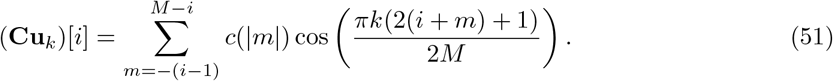

Use the Trigonometric identity:

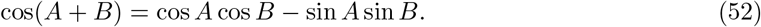

Set 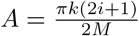 and 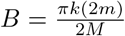, so:

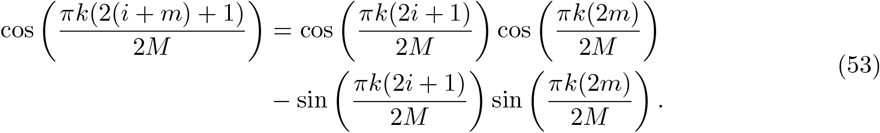

Substitute this back:

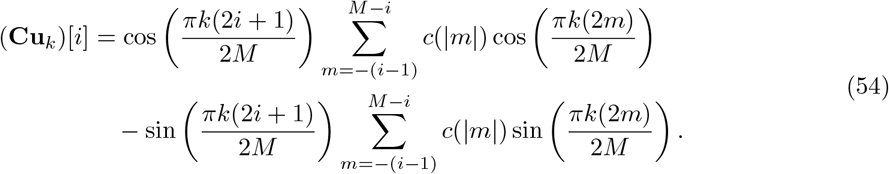

Since *c*(|*m*|) is an even function:

- 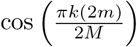 is even, so the sum involving it is nonzero.
- 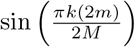 is odd, so the sum involving it is zero:

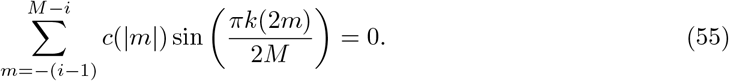

Thus:

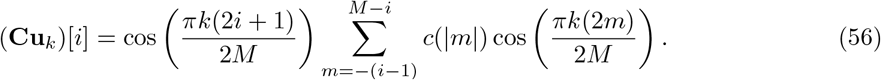

Assume *M* is large and *i* is not near the edges. Approximate the sum over all *m* from *−∞* to *∞*:

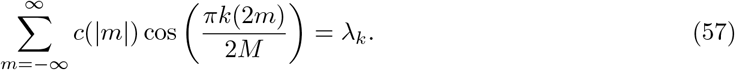

Let:

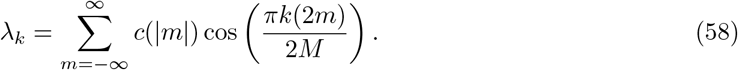

We have:

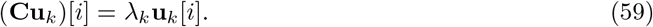

This shows that **u**_*k*_ is an eigenvector of **C**, with eigenvalue *λ*_*k*_.

## References

Bernardi, S., M.K. Benna, M. Rigotti, J. Munuera, S. Fusi, and C.D. Salzman. 2020. The geometry of abstraction in the hippocampus and prefrontal cortex. Cell 183 (4): 954–967.

Bottini, R. and C.F. Doeller. 2020. Knowledge across reference frames: Cognitive maps and image spaces. Trends in Cognitive Sciences 24 (8): 606–619.

Britanak, V., P.C. Yip, and K.R. Rao. 2010. Discrete cosine and sine transforms: general properties, fast algorithms and integer approximations. Elsevier.

Brunamonti, E., V. Mione, F. Di Bello, P. Pani, A. Genovesio, and S. Ferraina. 2016. Neuronal modulation in the prefrontal cortex in a transitive inference task: Evidence of neuronal correlates of mental schema management. Journal of Neuroscience 36 (4): 1223–1236.

Chafee, M.V. 2013. A scalar neural code for categories in parietal cortex: Representing cognitive variables as “more” or “less”. Neuron 77 (1): 7–9.

Chéreau, R., L.E. Williams, T. Bawa, and A. Holtmaat 2022. Circuit mechanisms for cortical plasticity and learning. In Seminars in cell & developmental biology, Volume 125, pp. 68–75. Elsevier.

Courellis, H.S., J. Mixha, A.R. Cardenas, D. Kimmel, C.M. Reed, T.A. Valiante, C.D. Salzman, A.N. Mamelak, R. Adolphs, S. Fusi, et al. 2023. Abstract representations emerge in human hippocampal neurons during inference behavior. bioRxiv : 2023–11.

De Falco, E., M.J. Ison, I. Fried, and R. Quian Quiroga. 2016. Long-term coding of personal and universal associations underlying the memory web in the human brain. Nature communications 7 (1): 13408.

Di Antonio, G., S. Raglio, and M. Mattia. 2024. A geometrical solution underlies general neural principle for serial ordering. Nature Communications 15 (1): 8238.

Fitzgerald, J.K., D.J. Freedman, A. Fanini, S. Bennur, J.I. Gold, and J.A. Assad. 2013. Biased associative representations in parietal cortex. Neuron 77 (1): 180–191.

Fried, I. 2022. Neurons as will and representation. Nature Reviews Neuroscience 23 (2): 104–114.

Ganguli, S., J.W. Bisley, J.D. Roitman, M.N. Shadlen, M.E. Goldberg, and K.D. Miller. 2008. One-dimensional dynamics of attention and decision making in lip. Neuron 58 (1): 15–25.

Gardner, R.J., E. Hermansen, M. Pachitariu, Y. Burak, N.A. Baas, B.A. Dunn, M.B. Moser, and E.I. Moser. 2022. Toroidal topology of population activity in grid cells. Nature 602 (7895): 123–128.

Gray, R.M. et al. 2006. Toeplitz and circulant matrices: A review. Foundations and Trends® in Communications and Information Theory 2 (3): 155–239.

Heald, J.B., D.M. Wolpert, and M. Lengyel. 2023. The computational and neural bases of context-dependent learning. Annual Review of Neuroscience 46 (1): 233–258.

Hong, S.Z., L. Mesik, C.D. Grossman, J.Y. Cohen, B. Lee, D. Severin, H.K. Lee, J.W. Hell, and A. Kirkwood. 2022. Norepinephrine potentiates and serotonin depresses visual cortical responses by transforming eligibility traces. Nature communications 13 (1): 3202.

Kumano, H., Y. Suda, and T. Uka. 2016. Context-dependent accumulation of sensory evidence in the parietal cortex underlies flexible task switching. Journal of Neuroscience 36 (48): 12192–12202.

Li, C., Y. Li, H. Jiang, W. Song, P. Lin, Z. Wang, J.J. Yang, Q. Xia, M. Hu, E. Montgomery, et al. 2018. Large memristor crossbars for analog computing. In 2018 IEEE International Symposium on Circuits and Systems (ISCAS), pp. 1–4. IEEE.

Lippl, S., K. Kay, G. Jensen, V.P. Ferrera, and L. Abbott. 2024. A mathematical theory of relational generalization in transitive inference. Proceedings of the National Academy of Sciences 121 (28): e2314511121.

Lorenzi, E., M. Perrino, and G. Vallortigara. 2021. Numerosities and other magnitudes in the brains: a comparative view. Frontiers in Psychology 12: 641994.

Luyckx, F., H. Nili, B. Spitzer, and C. Summerfield. 2019. Neural structure mapping in human probabilistic reward learning. elife 8: e42816.

Lyons, I.M., S.E. Vogel, and D. Ansari. 2016. On the ordinality of numbers: A review of neural and behavioral studies. Progress in brain research 227: 187–221.

Magee, J.C. and C. Grienberger. 2020. Synaptic plasticity forms and functions. Annual review of neuroscience 43 (1): 95–117.

Mannella, F. and G. Pezzulo. 2024. Transitive inference as probabilistic preference learning. Psychonomic Bulletin & Review : 1–16.

Markram, H., J. Lübke, M. Frotscher, and B. Sakmann. 1997. Regulation of synaptic efficacy by coincidence of postsynaptic aps and epsps. Science 275 (5297): 213–215.

Merritt, D.J. and H.S. Terrace. 2011. Mechanisms of inferential order judgments in humans (homo sapiens) and rhesus monkeys (macaca mulatta). Journal of Comparative Psychology 125 (2): 227.

Miconi, T. and K. Kay. 2025. Neural mechanisms of relational learning and fast knowledge reassembly in plastic neural networks. Nature Neuroscience: 1–9.

Moyer, R.S. and R.H. Bayer. 1976. Mental comparison and the symbolic distance effect. Cognitive psychology 8 (2): 228–246.

Nelli, S., L. Braun, T. Dumbalska, A. Saxe, and C. Summerfield. 2023. Neural knowledge assembly in humans and neural networks. Neuron 111 (9): 1504–1516.

Nieder, A. 2016. The neuronal code for number. Nature Reviews Neuroscience 17 (6): 366–382.

Nieder, A. and S. Dehaene. 2009. Representation of number in the brain. Annual review of neuroscience 32 (1): 185–208.

Okazawa, G., C.E. Hatch, A. Mancoo, C.K. Machens, and R. Kiani. 2021. Representational geometry of perceptual decisions in the monkey parietal cortex. Cell 184 (14): 3748–3761.

Roitman, J.D., E.M. Brannon, and M.L. Platt. 2012. Representation of numerosity in posterior parietal cortex. Frontiers in Integrative Neuroscience 6: 25.

Rutishauser, U., T. Aflalo, E.R. Rosario, N. Pouratian, and R.A. Andersen. 2018. Single-neuron representation of memory strength and recognition confidence in left human posterior parietal cortex. Neuron 97 (1): 209–220.

Sorscher, B., G.C. Mel, S.A. Ocko, L.M. Giocomo, and S. Ganguli. 2023. A unified theory for the computational and mechanistic origins of grid cells. Neuron 111 (1): 121–137.

Summerfield, C., F. Luyckx, and H. Sheahan. 2020. Structure learning and the posterior parietal cortex. Progress in neurobiology 184: 101717.

Sun, W., J. Winnubst, M. Natrajan, C. Lai, K. Kajikawa, A. Bast, M. Michaelos, R. Gattoni, C. Stringer, D. Flickinger, et al. 2025. Learning produces an orthogonalized state machine in the hippocampus. Nature: 1–11.

Viganò, S., R. Bayramova, C.F. Doeller, and R. Bottini. 2024. Spontaneous eye movements reflect the representational geometries of conceptual spaces. Proceedings of the National Academy of Sciences 121 (17): e2403858121.

Waydo, S., A. Kraskov, R.Q. Quiroga, I. Fried, and C. Koch. 2006. Sparse representation in the human medial temporal lobe. Journal of Neuroscience 26 (40): 10232–10234.

Xiao, Z., X. Wang, J. Zhang, J. Ou, L. He, Y. Qu, X. Hu, T. Behrens, and Y. Liu. 2025. Human hippocampal ripples predict the alignment of experience to a grid-like schema. bioRxiv : 2025–01.

Xu, B., J. Wu, H. Xiao, T.F. Münte, and Z. Ye. 2024. Inferior parietal cortex represents relational structures for explicit transitive inference. Cerebral Cortex 34 (4): bhae137.

